# *In vitro* Pharmacological Evaluation, *In silico* pharmacokinetics and TOPKAT validation of *Eichhornia crassipes*: Formulation of a Collagen-Based Matrix for Wound Healing Applications

**DOI:** 10.1101/2025.08.25.672252

**Authors:** S. Venkata Krishnan, C. Shanmuga Sundaram

**Affiliations:** HINDUSTAN COLLEGE OF ARTS & SCIENCE

**Keywords:** Eichhornia crassipe, Membrane Stabilization, Molecular docking, TOPKAT

## Abstract

The aspiration of this study was to develop a high target potency drug to aggrandize the normal wound healing progresses through inflammatory, proliferative and remodeling phases in response to tissue injury. One of the essential components of the extracellular matrix is collagen, which serves as indispensable for controlling the various stages of wound healing. The *Eichhornia crassipes* extract exhibited significant free radical scavenging activities with 72.1% and 80% inhibition in DPPH and ABTS assays at 0.1 mg/ml, respectively. Notably, it showed 80.2% inhibition of α-amylase and 87.3% of α-glucosidase, indicating strong antidiabetic potential. In anti-inflammatory assays, the extract demonstrated effective protection against RBC lysis, highlighting its membrane-stabilizing properties. Antibacterial tests revealed notable activity against various pathogens, with inhibition zones up to 15 mm. Cytotoxicity assays indicated variable effects on Vero cell morphology, while wound healing studies showed enhanced cell migration and scratch closure rates. The results were then validated by the use of Molecular docking, oral bioavailability parameters (TOPKAT), ADMET risk screening and *in-silico* pharmacokinetics and pharmacodynamics analysis which identified phytochemicals indicated promising drug-like properties.

## Introduction

The skin serves as a protective barrier against pathogen invasion. Wounds occur when the normal anatomical structure was disrupted, whether through surgical procedures or due to chemical, physical, mechanical and thermal injuries, all of which also affect skin function (Maillard et al., 2021). Because the skin was frequently exposed to external elements, such as wounds, scrapes and environmental interactions, it is more susceptible to pathogen colonization. Wounds can be classified into two types: acute and chronic. Acute wounds, caused by external factors, heal through the typical stages of wound repair, including cuts, burns, abrasions and surgical wounds (Pallavali et al., 2017). In contrast, chronic wounds are characterized by persistent infections, necrosis and ongoing inflammation, preventing the wound from healing (Raziyeva et al., 2021). A study by Ashoobi et al. (2023) analyzed 506 patients and found surgical wound infections in 24 cases (4.7%), while non-surgical wound infections had a higher rate of developing into surgical wound infections (12.5%). Wound infections are responsible for 70–80% of deaths in surgical patients and account for one-third of nosocomial infections. Infected wounds significantly impact quality of life and slow the healing process. Regardless of the wound type, infections are associated with increased patient morbidity and mortality, particularly in developing countries (Mehta et al., 2007). Chronic wounds or surgical wound infections that fail to heal lead to higher healthcare costs due to longer hospital stays, extensive diagnostic tests, prolonged antibiotic treatments and sometimes invasive surgeries (Puca et al., 2019). Effective wound care and antibiotic therapy are crucial for managing infections (Jeschke et al., 2020). On the other hand, chronic wounds, such as leg or arterial ulcers, heal more slowly and may result from internal conditions like immunological deficiencies or diabetes (Vaez and Beigi, 2016). Various dressing materials have been developed for wound treatment and care, with a focus on enhancing the healing process and preventing microbial infections (Dhivya et al., 2015). Biological materials, such as proteins (keratin, fibrin, collagen, etc.) and polysaccharides (chitosan, alginate, fucoidan, etc.), have gained popularity in recent years for use as wound dressings due to their biocompatibility, biodegradability and non-toxicity (Murray et al., 2019). Collagens, the most common structural proteins, are found in connective tissues, skin and bone, forming fibrils with a triple helical structure. Among the 28 types of collagen, type I collagen was the most abundant (Pallaske et al., 2018). Other notable types, such as types II, III and IV, are found in fish tissues. Like type I collagen, type II collagen also forms fibrils but has a different polypeptide chain composition (Ricard-Blum, 2011). Type II collagen, which is predominant in cartilage, plays a key role in providing tensile strength to the tissue (Eyre, 2001). Although type II collagen derived from marine sources has been widely used in bone and cartilage tissue engineering and regeneration, its application in acute skin wound healing was less explored (Lai et al., 2020).

Water hyacinth, scientifically known as *Eichhornia crassipes* (Mart.), is a free-floating, monocotyledonous aquatic plant from the Pontederiaceae family. Originally native to Brazil and the Amazon, it has since spread to tropical and subtropical regions worldwide. The plant was introduced to India by the British at the end of the 18th century, after the British Governor-General was captivated by its flowers and brought it with him. Water hyacinth has since proliferated in many water bodies across India. It has also been reported in numerous African countries, including Egypt, Sudan, Kenya, Ethiopia, Nigeria, Zimbabwe, Zambia and South Africa (Dersseh et al., 2019). The plant is known for its remarkable ability to tolerate temperature variations, changes in pH and nutrient levels and its rapid spread. Consequently, it was ranked among the 10 worst weed plants in the world and one of the 100 most aggressive invasive species by the International Union for Conservation of Nature (Téllez et al., 2008). In Nigeria, water hyacinth was used for skin treatments and a mixture of rice flour, turmeric and plant leaf extract has been used to treat eczema, attributed to the plant’s high vitamin C content (Sharma et al., 2020). The stems and leaves of *E. crassipes* are also used for their anti-inflammatory properties, which are linked to the plant’s phenolic compounds, to treat wounds and swelling (Rorong et al., 2012). In the Philippines, a traditional topical anti-inflammatory remedy was made from lemon juice combined with the juice of *E. crassipes* leaves (Sharma et al., 2020). The current study aims at the potential of plant for wound healing and its underlying molecular mechanisms are currently being explored for possible applications in the cosmeceutical industry (Bakrim et al., 2022).

## Material and Methods

### Sample collection and processing

The plant sample *Eichhornia crassipes* was collected from Sithalapakkam Lake (12.8865° N, 80.1777° E), Chennai, Tamil Nadu, India. The sample was authenticated by Captain Srinivasa Murthy Central Ayurveda Research Institute, Central Council for Research in Ayurvedic Sciences Ministry of AYUSH, Chennai, Tamil Nadu, India. The plant material was thoroughly cleaned, cut into small pieces and shade-dried at room temperature. Once dried, the samples were ground into a coarse powder (70–80% yield) and stored in airtight containers for extraction.

### Preparation of ethanol extract

The powdered sample of *E. crassipes* (50 g) was extracted with ethanol using a Soxhlet apparatus for 24 hours (∼20 cycles) at a temperature of 75–80^°^C. The resulting extract was then concentrated and dried on a water bath at 50^°^C to obtain a semisolid mass. The concentrated extract was stored in an airtight container for further use.

### Antioxidant activity

The antioxidant activity of the sample was evaluated using the 1,1-diphenyl-2-picrylhydrazyl (DPPH) method (Blois, 1958) and the 2,2’-azinobis-(3-ethylbenzothiazoline-6-sulfonic acid) (ABTS) method (Shirwaikar et al., 2006). For the DPPH assay, a 0.4 mM DPPH solution in methanol was prepared. For the ABTS assay, an ABTS solution (7 mM ABTS and 100 mM potassium persulfate) in methanol was prepared. Two milliliters of each solution were added to different concentrations (20, 40, 60, 80, 100 µg/ml) of the ethanol extract of *E. crassipes* leaf, as Benchmark control 5 µg/ml Ascorbic Acid was prepared and added Independently as same concentration of plant extract. The mixtures were then allowed to stand in the dark at room temperature for 15 minutes. Absorbance was measured at 517 nm for DPPH and at 734 nm for ABTS, using methanol as a blank. A lower absorbance indicated a higher free radical scavenging activity. The percentage of radical scavenging was calculated using the following equation: Radical scavenging activity (%) = ((Absorbance control - Absorbance sample) /Absorbance control) x 100.

### *In vitro* antidiabetic activity - α-amylase inhibitory activity

The inhibitory effect of the ethanol extract of *E. crassipes* leaf on α-amylase activity was evaluated following the protocol by Thalapeneni et al. (2008). Various concentrations (20, 40, 60, 80, 100 µg/ml) of the ethanol leaf extract and as Benchmark control 5 µg/ml Metformin was prepared and added consecutively as same concentration of plant extract were mixed with 500 µl of porcine pancreatic α-amylase (0.5 mg/ml, prepared in 0.20 mM phosphate buffer at pH 6.9) solution separately. Next, 500 µl of 1% starch solution was introduced and the mixture was incubated for 10 minutes at 25°C. The reaction was stopped by adding DNS (3,5-dinitrosalicylic acid reagent) and the test tubes were placed in a boiling water bath for 5 minutes, then cooled and diluted with 10000 µl of Milli-Q water. The absorbance was measured at 540 nm. The percentage inhibition of α-amylase by the plant extract was calculated as follows: Inhibition (%) = (Abs control - Abs sample /Abs control) x 100.

### *In vitro* antidiabetic activity - α-glucosidase inhibitory activity

The α-glucosidase inhibitory activity of the *E. crassipes* leaf extract was evaluated following the method of Andrade-Cetto et al. (2008). Different concentrations (20, 40, 60, 80 and 100 µg/ml) of the ethanolic leaf extract and as Benchmark control 5 µg/ml Metformin was prepared and added in a distinct reaction mixture as same concentration of plant extract were incubated with 500 µl of phosphate buffer (100 mM, pH 6.8) and 250 µl of p-nitrophenyl-α-D-glucopyranoside solution (5 mM) in phosphate buffer (100 mM, pH 6.8) at 37°C for 15 minutes. Subsequently, 250 µl of α-glucosidase enzyme solution (1.5 U/ml) was added and the mixture was incubated at 37°C for another 15 minutes. The reaction was stopped by adding 2000 µl of 200 mM sodium carbonate solution. The absorbance of the released p-nitrophenol was measured at 405 nm. The inhibitory activity was calculated as a percentage relative to the control (without inhibitors) using the following formula, Inhibition (%) = (Abs sample /Abs control) - 1) x 100.

### Preparation of collagen-*E. crassipes* extract matrix

The matrix was prepared following the protocol of Gaspar-Pintiliescu et al. (2018) with slight modifications. A collagen solution was first made by dissolving 0.5 g of type-I collagen (HiMedia) in 50000 µl of double-distilled water at pH 5.5. Collagen (Col) and *E. crassipes* ethanol extract (ECEE) matrices were then prepared in different ratios (3:1, 5:1 and 10:1 v/v) by stirring the mixtures for 2 hours using a magnetic stirrer. The resulting Col-ECEE matrices were stored in airtight containers for further analysis.

### *In vitro* anti-inflammatory activity

The anti-inflammatory activity of the *E. crassipes* leaf extract and Col-ECEE matrices was evaluated using the *In vitro* HRBC (human red blood cell) membrane stabilization method (Chowdhury et al., 2014). Blood was collected from healthy volunteers and mixed with an equal volume of Alsever solution (containing 2% dextrose, 0.8% sodium citrate, 0.05% citric acid and 0.42% sodium chloride in 100 ml distilled water) and 500 µl of isosaline, then centrifuged at 2500 rpm. To 1000 µl of HRBC suspension, an equal volume of the test drug at three concentrations (100, 200 and 300 µg/ml) was added. The assay mixtures were incubated at 37°C for 30 minutes and then centrifuged at 5000 rpm. The hemoglobin content in the supernatant was measured using a spectrophotometer at 560 nm. The percentage of hemolysis was calculated using the formula, Hemolysis (%) = (Absorbance of Test /Absorbance of Control) × 100. The percentage of protection was then calculated as, Protection (%) = 100 - Hemolysis (%).

### Antibacterial activity - Agar well diffusion method

The antibacterial activity of the samples was evaluated using the well diffusion method (Balouiri et al., 2016). Fresh bacterial cultures of the target pathogens, such as *Escherichia coli, Pseudomonas aeruginosa, Bacillus subtilis* and *Staphylococcus aureus*, were prepared by inoculating them in nutrient broth and incubating at 37°C for 18-24 hours until they reached the desired turbidity (usually 0.5 McFarland standard). A uniform lawn of the bacterial culture was swabbed on the nutrient agar plates and wells of 5 mm in diameter were created using a sterile cork borer, ensuring the wells were evenly spaced. The test samples, *E. crassipes* leaf extract, collagen alone and 5:1 Col-ECEE matrix were added at concentration of 50 µg/ml into separate wells. A control well containing 20 µl gentamycin (10 µg) was also included as a positive drug. The plates were incubated at 37°C for 24 hours under aerobic conditions. After incubation, the antibacterial activity was assessed by measuring the diameter of the clear inhibition zones around each well (in mm). Larger inhibition zones indicated stronger antibacterial activity.

### Cytotoxic effect

Vero (African green monkey kidney normal epithelial cell line) was obtained from the National Centre for Cell Sciences (NCCS), Pune, India. Cells were maintained in the logarithmic phase of growth in Dulbecco’s modified eagle medium (DMEM) supplemented with 10% (v/v) heat inactivated fetal bovine serum (FBS), 100 U/ml penicillin, 100 μg/ml streptomycin. They were maintained at 37°C with 5% CO2 in 95% air humidified incubator. The cytotoxic effect of the samples was tested against Vero cell line by MTT (3-(4,5-dimethylthiazol-2-yl)-2,5-diphenyltetrazolium bromide assay (Mossman, 1983). Vero cells were seeded in 96-well microplates (1 x 10^4^ cells/well) and incubated at 37°C for 48 h to reach 70-80% confluence. Then the cells were treated with different concentrations (20, 40, 60, 80 and 100 µg/ml) of *E. crassipes* leaf extract, collagen alone and 5:1 Col-ECEE matrix and incubated for 24 h. The morphological changes of untreated (control) and the treated cells were observed under digital inverted microscope (20X magnification) after 24 h and photographed. The cells were then washed with phosphate-buffer saline (PBS, pH-7.4) and 20 μl of (MTT) solution (5 mg/ml in PBS) was added to each well. The plates were then stand at 37ºC in the dark for 4 h. The formazan crystals were dissolved in 100 μl DMSO and the absorbance was read spectrometrically at 570 nm. Percentage of cell viability was calculated using the formula, Cell viability (%) = (Absorbance of sample/Absorbance of control) X 100.

### *In vitro* scratch wound healing assay

*In vitro* scratch wound healing assay was performed according to the previously reported and standardized protocol (Liang et al., 2007). Vero cell were seeded in 6-well plates (1 × 10^5^ cells/well) and grown until reached a confluence of 90%, in the optimum culture conditions. In the middle of cell monolayer, a scratch was made by a P10 pipette tip, to mimic a wound and cell debris were removed by washing with fresh medium. The wound was exposed with 5:1 Col-ECEE matrix (50 µg/ml) for 24-48 h at 37°C in a humidified atmosphere of 5 % CO2. The negative control cells were maintained without any treatment. Scratch wound closure was analyzed in two modalities (Control and 5:1 Col-ECEE matrix) under the digital inverted microscope, by acquiring digital images at time 0 (T0), 24 hrs (T1) and 48 hrs (T2) (static imaging). The closure of the scratch was quantified by measuring the difference between the wound widths at T0 and T1/T2, using the ImageJ processing software and final image clarity and labelling for figures were enhanced using Canva (https://www.canva.com). Scratch closure rate (SCR) was calculated as described by Felice et al., (2015), using the following formula: SCR = ((T0 – T1/T2) /T0) × 100, Where T0 is the scratch area at time 0 and T1/T2 is the scratch area at 24 hrs and 48 hrs.

### Spectroscopic studies and imaging

The *E. crassipes* leaf extract and 5:1 Col-ECEE matrix were analyzed for the functional groups in the region of 4000 −400 cm-1 wave numbers at 4 cm-1 resolution by ATR-FTIR spectrometer (Thermo Nicolet Avatar 370 spectrometer). The microstructure of the 5:1 Col-ECEE matrix was observed under a scanning electron microscopy (SEM, JEOL JSM-6390LV). The *E. crassipes* leaf extract was analyzed by GC-MS using the Agilent 7890B Gas Chromatograph coupled with a 5977A Mass Spectrometer.

### Molecular docking and drug likeness analysis

Molecular docking studies were performed using the Discovery Studio Client software with the LibDock scoring function to evaluate the binding affinity between the ligand and the receptor. A higher score indicates a stronger receptor-ligand binding affinity (Daina et al., 2017). The scoring functions were utilized to assess ligand-binding affinities and screen out active and inactive compounds during virtual screening. The 3D structures of the phytochemicals were obtained from the PubChem database and used as input for docking (Singh et al., 2016). The docking targets included ‘7WEU’ (Crystal structure of Peroxiredoxin I), ‘3BAJ’ (Human Pancreatic Alpha-Amylase), ‘2QLY’ (Crystal Structure of the N-terminal Subunit of Human Maltase-Glucoamylase) and ‘3KBX’ (Human macrophage inflammatory protein-1 alpha L3M_V63M). The docking region was defined by removing any pre-existing ligand from the PDB file or receptor cavities, followed by re-docking the protein. The 3D docked protein-ligand complex, 2D interaction images, receptor surface interaction, AlogP98 and 2D polar surface area were analyzed using the Discovery Studio Client software to identify the interacting amino acid residues in the protein. Drug-like properties, based on Lipinski’s Rule of Five and bioactivity scores, were assessed using the Molinspiration webserver (www.molinspiration.com). ADME (Absorption, Distribution, Metabolism, Excretion) properties of the molecules were evaluated using SWISSADME (www.swissadme.ch).

## Results and discussions

The ethanolic leaf extract of *E. crassipes* was evaluated for its scavenging potential against DPPH and ABTS free radicals. Lee et al. (2015) confirmed that compounds with high hydrogen-donating ability are effective in eliminating hydroxyl or superoxide radicals, either through physiological action or oxidation. At the highest concentration of 100 µg/ml, *E. crassipes* extract showed 72.1% inhibition in the DPPH assay and 80% inhibition in the ABTS assay (Fig. 1). The IC50 values were determined to be 21.35 µg/ml for the DPPH assay and 18.75 µg/ml for the ABTS assay. As a benchmark control, ascorbic acid at 5 µg/ml exhibited highest of 91.13% inhibition in the DPPH assay and 97.41% inhibition in the ABTS assay, confirming the reliability of the assay system (Fig. 2). Noufal et al. (2023) reported that the methanolic extract of *E. crassipes* exhibited antioxidant activities of 57.95% in DPPH and 60.47% in ABTS at a concentration of 400 µg/ml. According to Blois (1958), an inhibition percentage of ≥20% was considered significant for antioxidant activity. The relatively low IC_50_ values and high scavenging percentages observed with the *E. crassipes* extract suggest strong antioxidant potential, although slightly lower than the standard antioxidant ascorbic acid.

**Figure 1.**
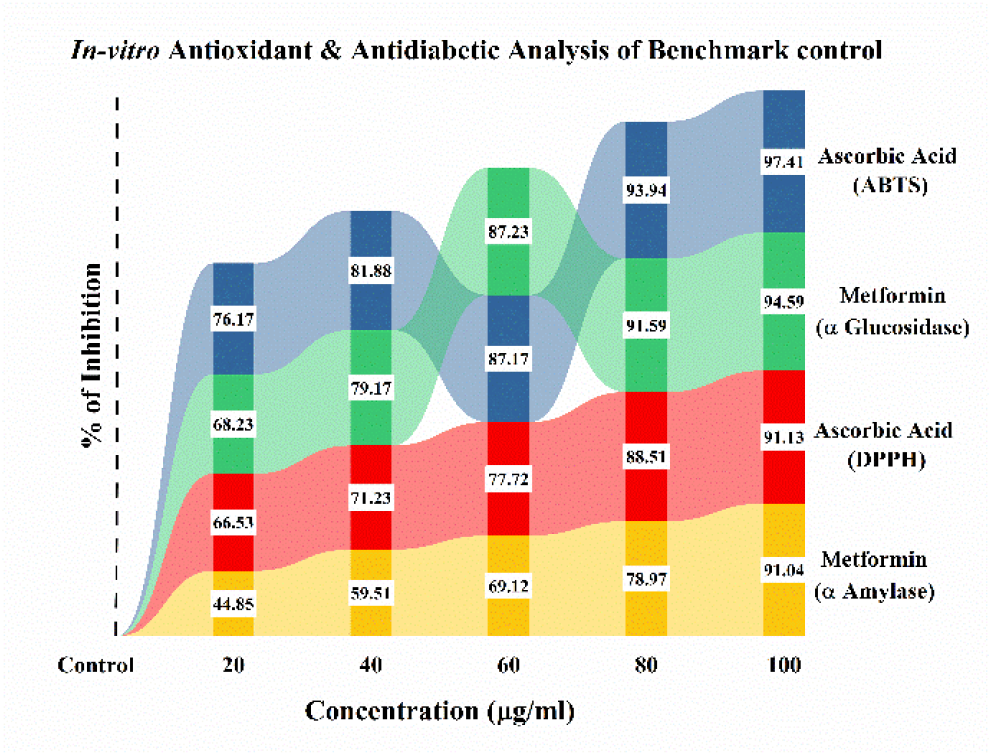
Ribbon chart of Antioxidant and *in vitro* antidiabetic analysis of Benchmark control

**Figure 2.**
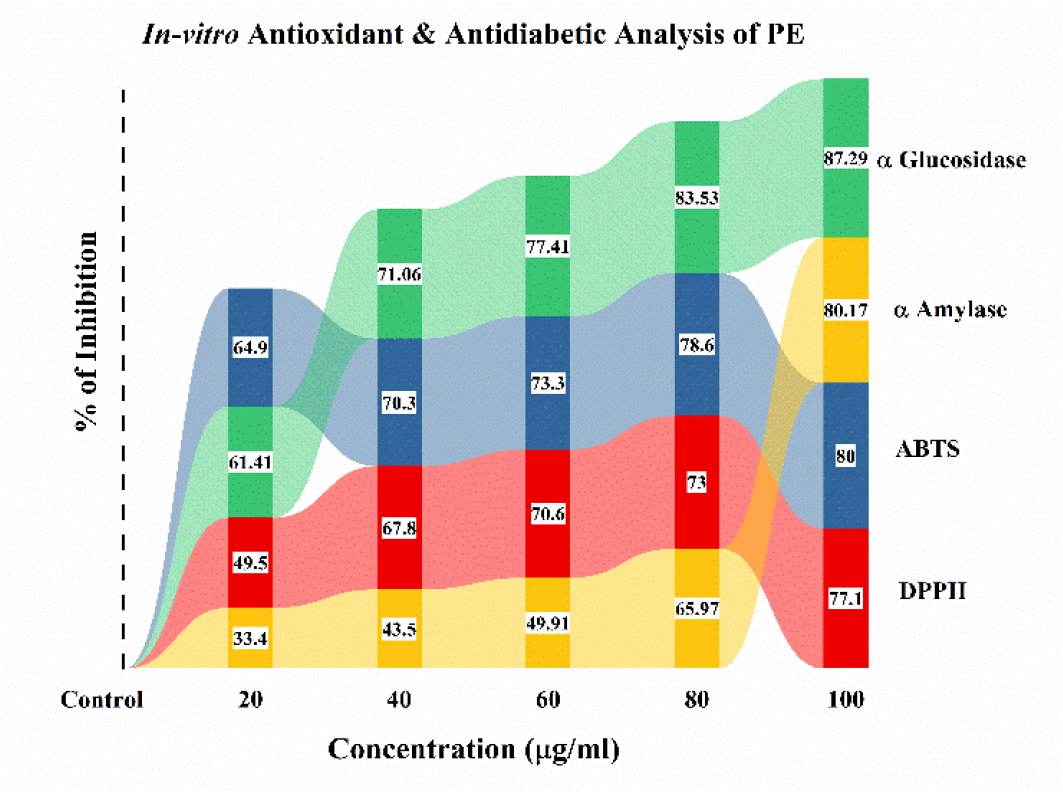
Ribbon chart of Antioxidant and *in vitro* antidiabetic analysis of ethanolic leaf extract of *E. crassipes*

Antidiabetic drugs are known to reduce blood glucose levels by inhibiting the activity of α-amylase and α-glucosidase enzymes. These enzymes, found in the brush border, play a crucial role in breaking down the by-products of luminal digestion into monosaccharides. α-Amylase is a calcium-dependent enzyme that facilitates starch digestion by converting it into glucose and maltose. Over-activity of this enzyme can contribute to postprandial hyperglycemia and by reducing starch digestion, blood glucose levels can be better managed (Kaur et al., 2021). Similarly, α-glucosidase, another brush border enzyme, hydrolyzes starches and disaccharides into glucose for absorption in the intestines. Inhibition of α-glucosidase delays carbohydrate absorption after meals, which helps lower postprandial glucose levels (Susan et al., 2016). The α-amylase inhibitory activity of *E. crassipes* leaf extract showed an 80.2% inhibition at 100 μg/ml, while the α-glucosidase inhibitory assay revealed a significant 87.3% inhibition at the same concentration (Fig. 1). The IC50 values was found to be 27.05 µg/ml and 24.59 µg/ml for α-amylase and α-glucosidase inhibition assays, respectively. As a benchmark control, Metformin at 5 µg/ml demonstrated 91.04% inhibition of α-amylase and 94.59% inhibition of α-glucosidase under the same experimental conditions, highlighting the potent enzyme inhibitory capacity of the standard antidiabetic drug (Fig 2). These results confirm the relevance and accuracy of the assay system, Olasunkanmi et al. (2021) reported that extracts from the roots, stems and leaves of *E. crassipes* exhibited significant inhibition, especially due to the presence of compounds like alkaloids, flavonoids, phenols, tannins, steroids and cardiac glycosides. The enzyme inhibitory effects of the *E. crassipes* leaf extract slightly lower than Metformin, suggested promising antidiabetic potential that warrants further pharmacological investigation.

Anti-inflammatory drugs typically work by inhibiting the cyclooxygenase (COX) enzyme, which was responsible for converting arachidonic acid into prostaglandins (PGs). Prostaglandin G2 (PGG2) was converted to PGH2 by the enzyme’s peroxidation activity, a process linked to the formation of prolonged channels in membranes. These channels allow biological mediators to be produced, enabling arachidonic acid to detach from the membrane and transform into prostaglandins. Both acute and chronic inflammation are thought to be associated with the extracellular activity of these enzymes (Varadarasu et al., 2007). Anti-inflammatory drugs act either by stabilizing the lysosomal membrane or inhibiting the enzyme cyclooxygenase. Since HRBC membranes are similar to lysosomal membranes, the ability of a drug to prevent hypotonicity-induced HRBC membrane lysis was used as an indicator of its anti-inflammatory activity (Chowdhury et al., 2014). The *E. crassipes* leaf extract and Col-ECEE matrices in 3:1, 5:1 and 10:1 ratios at concentrations of 80 and 100 µg/mL protects human erythrocyte membranes from lysis induced by a hypotonic solution. At the concentration of 100 µg/mL, the extract and matrices showed the highest inhibition of RBC hemolysis. Among the tested concentrations, the 5:1 Col-ECEE matrix and *E. crassipes* extract exhibited the lowest hemolysis percentages, with values of 57.55% and 56.11%, respectively. The *E. crassipes* extract provided the strongest protection against HRBC lysis with 43.88% protection, followed by the 5:1 Col-ECEE matrix (42.44%). The 3:1 and 10:1 Col-ECEE matrices showed lower protection, with values of 14.38% and 16.9%, respectively. The 5:1 Col-ECEE matrix was considered optimal, providing protection comparable to aspirin (52.15%) (Fig. 3).The results obtained indicate that RBC hemolysis was inhibited by dose-dependent manner of both the extract and matrices. It was well-established that a cell’s ability to remain viable depends on the integrity of its membranes (Haris et al., 1962). When red blood cells are exposed to harmful substances such as hypotonic media, methyl salicylate, or phenyl hydrazine, membrane lysis, hemolysis and oxidation of hemoglobin can occur (Ferrali et al., 1992). The hemolytic effect of hypotonic solutions was caused by an excessive influx of fluid into the cell, leading to membrane rupture. This damage to the RBC membrane increases the cell’s susceptibility to lipid peroxidation induced by free radicals, which can result in further cellular damage. This was consistent with the finding that the breakdown of biomolecules generates free radicals, exacerbating cellular damage (Maxwell, 1995). The accumulation of fluid in body tissues leads to edema in two phases. The first phase was induced by carrageenan, which releases kinins, serotonin and histamine, while the second phase was more evident and involves the release of prostaglandin-like compounds. Therefore, the anti-inflammatory potential of *E. crassipes* may stem from its ability to limit the production of inflammatory mediators or stabilize cellular membranes. These findings suggest that both the 5:1 Col-ECEE matrix and the *E. crassipes* ethanol extract exhibit stabilization of HRBC membrane and anti-inflammatory properties.

**Figure 3.**
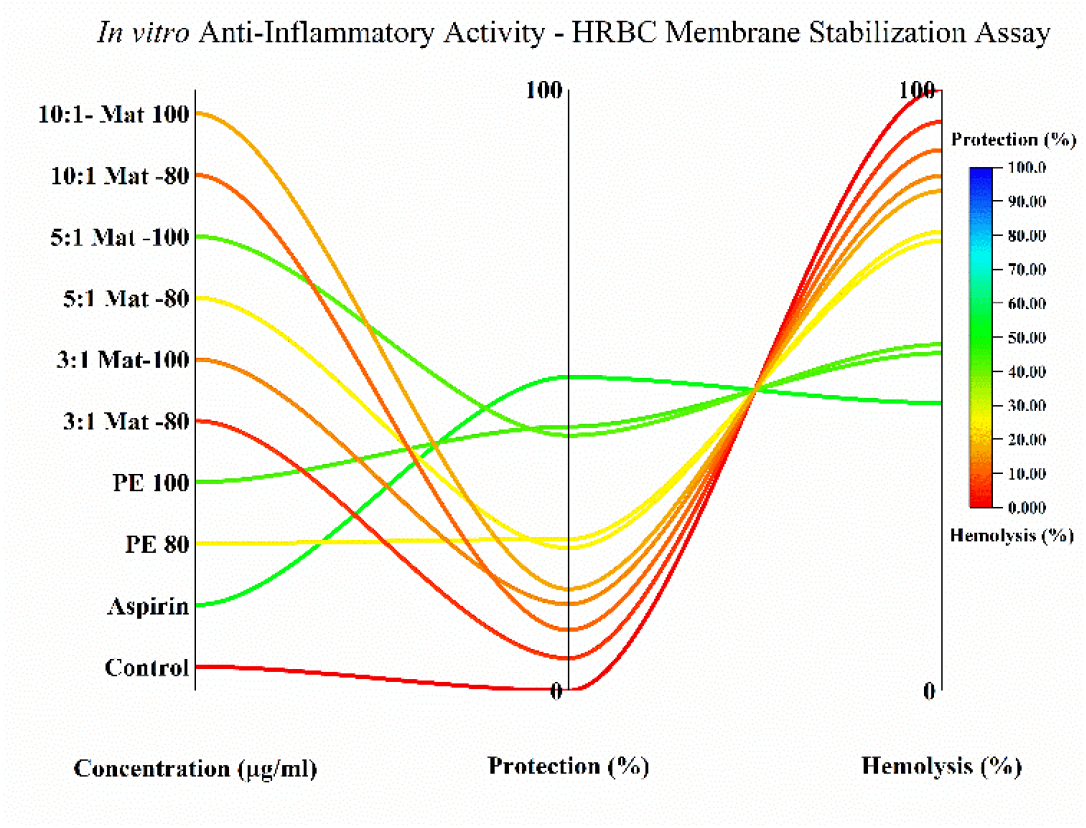
Parallel plot of *In vitro* anti-inflammatory activity

The antibacterial activity of the *E. crassipes* leaf extract, collagen alone and 5:1 Col-ECEE matrix was tested against *E. coli, P. aeruginosa, B. subtilis* and *S. aureus* showed varied responses as depicted in Fig. 4. The zones of inhibition observed were as follows: for *E. coli*, the plant extract showed a 12 mm zone and the 5:1 Col-ECEE matrix showed a 15 mm zone; for *B. subtilis*, the plant extract exhibited an 11 mm zone, while the 5:1 Col-ECEE matrix showed a 10 mm zone (Fig. 5). Both the plant extract and the 5:1 Col-ECEE matrix displayed similar inhibition zones of 12 mm against *S. aureus* and 10 mm against *P. aeruginosa*. Collagen alone did not show any antibacterial activity against the tested pathogens (Fig. 5). Zones of inhibition with a diameter of ≥10 mm are generally considered indicative of bactericidal activity (Usman et al., 2009). Haggag et al. (2016) reported that the N-butanol and methanol extracts of *E. crassipes* exhibited significant antibacterial and antifungal activities.

**Figure 4.**
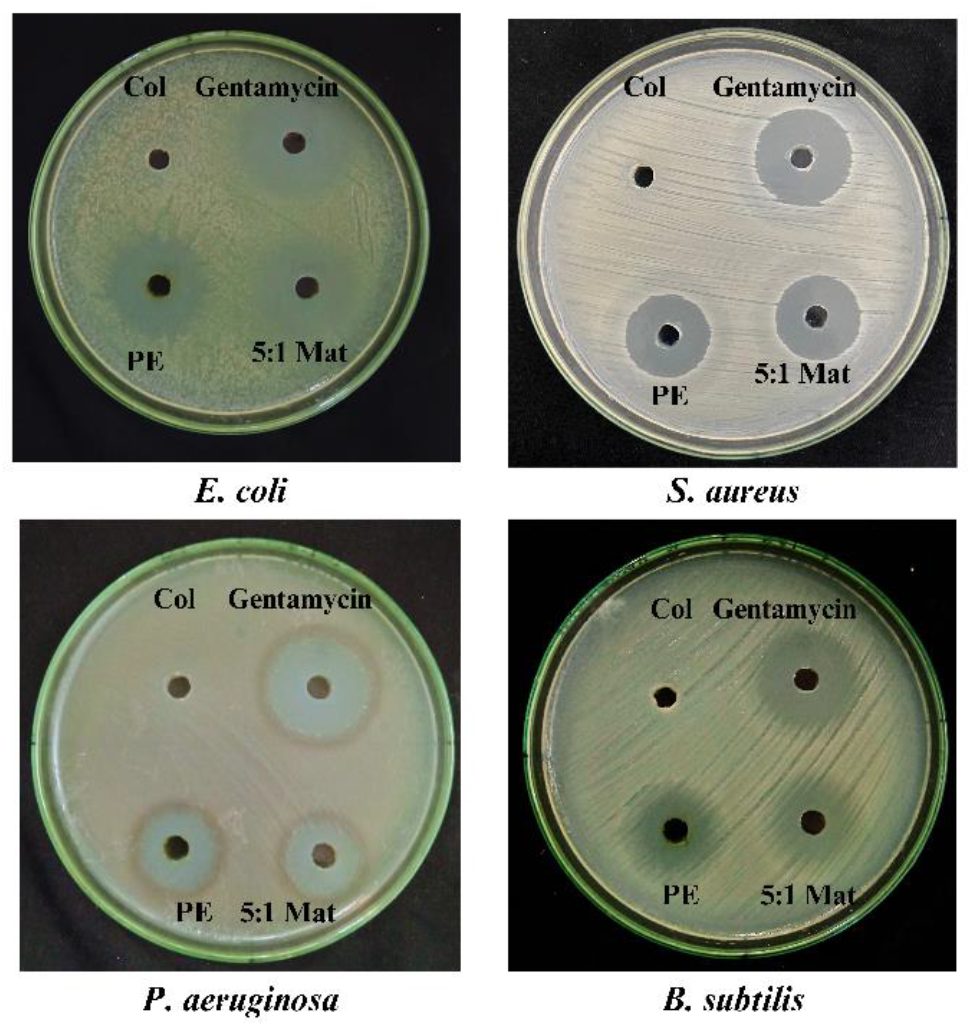
Antibacterial activity of *E. crassipes* leaf extract, collagen alone and 51 Col-ECEE matrix

**Figure 5.**
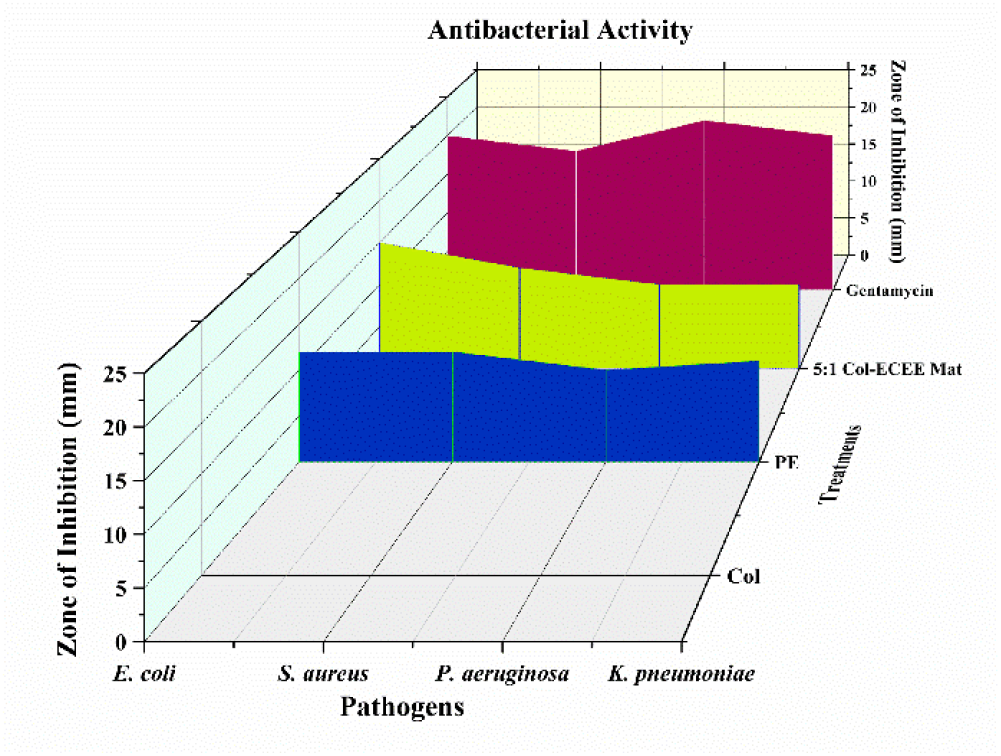
3D Waterfall Graph - Zone of inhibition measurements of *E. crassipes* leaf extract, collagen alone and 51 Col-ECEE matrix against tested pathogens

Cytotoxicity assays are valuable for understanding the toxicity of substances in terms of time- and dose-dependence, as well as their impact on the cell cycle and potential reversibility (Gupta et al., 2022). The cytotoxic effects of *E. crassipes* leaf extract, collagen alone and the 5:1 Col-ECEE matrix were assessed using the Vero cell line. The results indicated a dose-dependent response, with varying effects on the morphology and viability of the cells. The morphological changes in both untreated (control) and treated cells (at a concentration of 100 µg/ml) are shown in Fig. 6. In the control group, Vero cells maintained their characteristic morphology, displaying a typical spindle shape with elongated ends (Fig. 6). This confirmed the normal, healthy state of the cells, with no exposure to any toxic substances. The cells exhibited typical cellular architecture, indicative of normal growth and viability.

**Figure 6.**
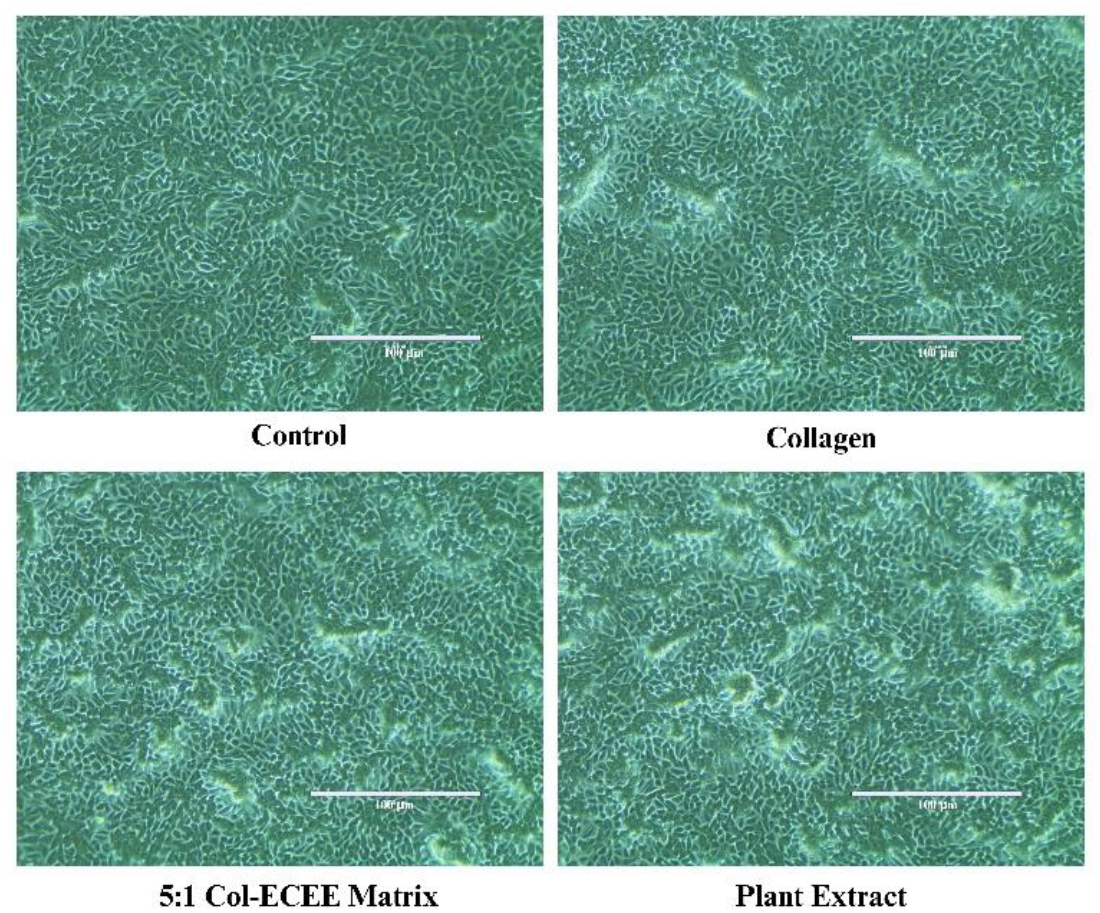
*In vitro* cytotoxicity activity of *E. crassipes* leaf extract, collagen alone and the 51 Col-ECEE matrix on Vero cell line

At the concentration of 100 µg/ml, collagen alone and 5:1 Col-ECEE matrix demonstrated a significant cell viability of 90.15% and 78.78% (Fig. 7). There were minimal changes in cell morphology, with the cells maintaining their spindle shape and displaying minimal signs of damage or detachment. The *E. crassipes* leaf extract, however, exhibited more pronounced cytotoxic effects at the same concentration (100 µg/ml), with several cells becoming round and detached from the culture surface (Fig. 6). The cell viability was found to be 63.98% which suggested that the extract may have a mild cytotoxic effect, which becomes more evident at higher concentrations. However, the combination of *E. crassipes* extract with collagen in the 5:1 Col-ECEE matrix helped to mask the extract’s cytotoxicity, suggesting that collagen may provide a protective or stabilizing effect on the cells.The 5:1 Col-ECEE matrix appears to be a promising material for biomedical applications, combining the biocompatibility of collagen with the potential therapeutic benefits of *E. crassipes* leaf extract, while minimizing toxicity. These results are consistent with findings in previous studies, which have shown that plant extracts can cause cell damage at higher concentrations (Uğur et al., 2017; Efenberger-Szmechtyk et al., 2020; Aldbass et al., 2021).

**Figure 7.**
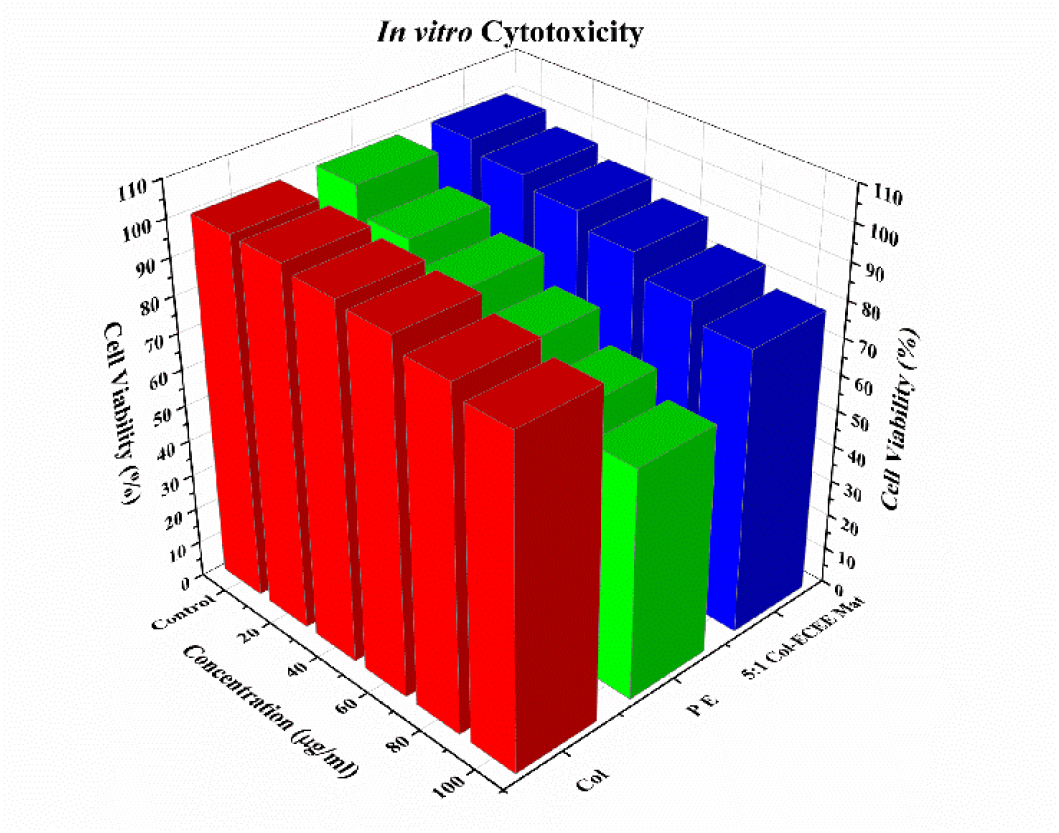
3D bar graph - Percentage of cell viability against Vero cell line

Fibroblast cell migration to the wound bed promotes the development of extracellular matrix (ECM) elements that are essential for the healing of wounds (Rousselle et al., 2019). Because of their chemotactic properties, low molecular weight peptides have been shown to promote wound healing cell migration (Tracy et al., 2016). It was well recognized that specific low molecular weight peptide motifs including the tripeptides arginine, glycine and asparagine are crucial for integrin-mediated cell adhesion and migration because they create recognition sites for integrins (cell membrane receptors) (Ruoslahti and Pierschbacher, 1987.) Proline-hydroxyproline dipeptide may play a role in the development of certain fibroblast mesenchymal stem cells, which are vital for wound healing. Therefore, low molecular weight peptides rich in proline and hydroxyproline amino acids found in collagen hydrolysate can serve as a great source for cell adhesion and proliferation (Manjushree et al., 2022). The *In vitro* wound healing activity of the 5:1 Col-ECEE matrix was evaluated using a standard scratch assay on Vero cells. This assay simulates the wound healing process by creating a “scratch” in a cell monolayer and the ability of the cells to migrate and close the wound was used to assess wound healing potential. The results indicate a significant effect of the 5:1 Col-ECEE matrix on promoting cell migration and wound closure, suggesting its potential as a wound healing agent. At the 50 µg/ml concentration of the 5:1 Col-ECEE matrix, the wound closure rate was significantly higher compared to the control group. After 24 hours of incubation, the scratch wound in the control group showed 38.18% migration of cells towards the centre of the wound, whereas, the cells treated with the 5:1 Col-ECEE matrix showed rapid migration and proliferation of 46.66%, with a noticeable reduction in the wound area (Fig. 8). The wound closure rate was also quantified at 48 hrs, revealing 71.36% and 91.42% closure in control and treated groups, respectively, indicating a robust wound healing potential of the matrix.

**Figure 8.**
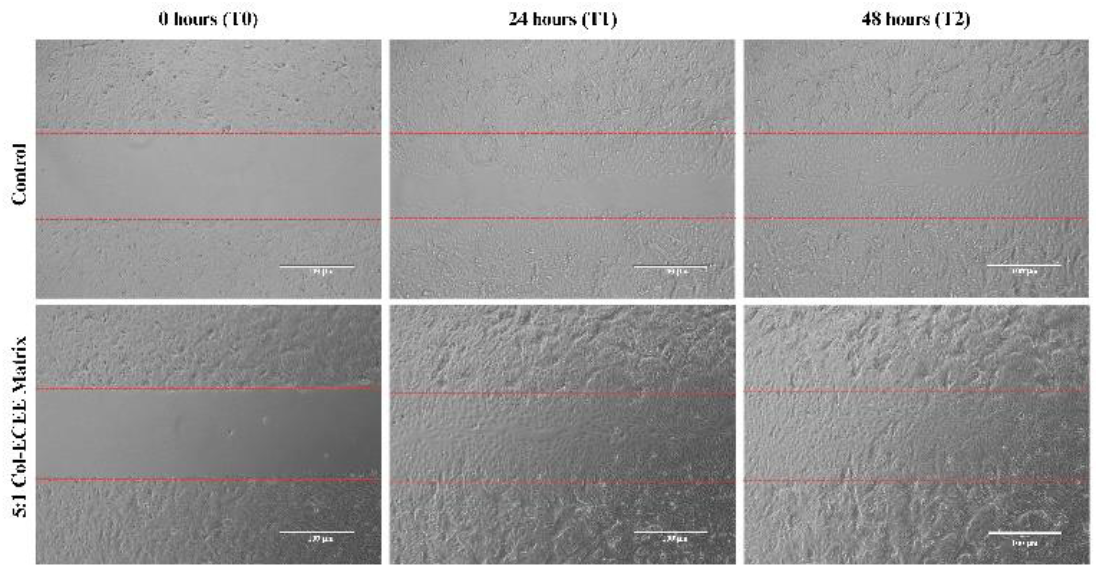
*In vitro* wound healing activity of 51 Col-ECEE matrix

The enhanced wound healing observed with the 5:1 Col-ECEE matrix can be attributed to the combined properties of collagen and the bioactive compounds present in the *E. crassipes* leaf extract. Collagen, being a natural extracellular matrix component, are known to promote cell adhesion, migration and proliferation, which are key processes in wound healing. It provides structural support to the cells, facilitating their movement towards the wound site and accelerating tissue regeneration (Mathew-Steiner et al., 2021). On the other hand, the *E. crassipes* leaf extract, known for its anti-inflammatory and antioxidant properties, likely plays a significant role in modulating the inflammatory response and reducing oxidative stress at the wound site. This combination of effects can accelerate the repair of damaged tissues and prevent excessive inflammation, which is often a barrier to efficient wound healing. The control group, which received no treatment, exhibited limited wound healing activity. This indicates that while cell migration and proliferation naturally occur, the absence of any therapeutic agent significantly slows the healing process. In comparison, the 5:1 Col-ECEE matrix demonstrated a significantly faster wound closure rate, highlighting its superior wound healing properties. Moreover, the matrix’s biocompatibility, as shown by the cytotoxicity assay, suggests that it can be used safely in wound healing applications without adverse effects on cell viability. This makes the 5:1 Col-ECEE matrix a promising candidate for the development of wound care products, including dressings, gels, or scaffolds for tissue engineering.

The FTIR spectra analysis compares the functional groups in *E. crassipes* leaf extract, collagen and the 5:1 Col-ECEE matrix, highlighting molecular interactions that indicate successful conjugation. The conjugation was evidenced by shifts and the appearance of new peaks in the FTIR spectra, suggesting enhanced stability and potential bioactivity in the matrix for biomedical applications. Fig 9 depicts the FTIR spectra of the ethanol extract of *E. crassipes* leaf (green), pure collagen (blue) and the 5:1 Col-ECEE matrix (red). In the *E. crassipes* extract spectrum, key peaks were observed at 3325.28 cm^−1^ (O-H stretching), 2972.31 cm^−1^ and 2883.58 cm^−1^ (C-H stretching), 1656.85 cm^−1^ (C=O stretching), 1413.82 cm^−1^ and 1381.03 cm^−1^ (C-H bending), 1327.03 cm^−1^ and 1274.95 cm^−1^ (C-N stretching) and 1087.85 cm^−1^ and 1045.42 cm^−1^ (C-O stretching), along with additional fingerprint region peaks like 879.54, 802.39, 613.36 and 432.05 cm^−1^, indicating complex phytochemical structures such as phenolics, amines and carbonyl compounds.

**Figure 9.**
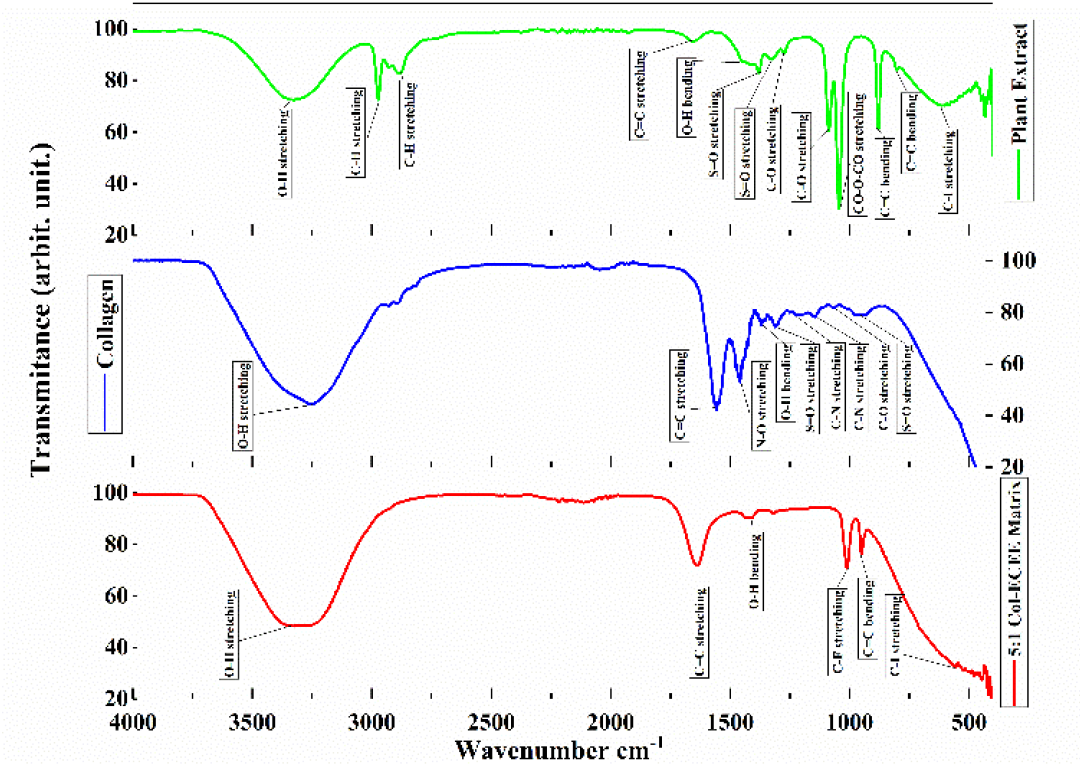
FTIR spectrum of (green*) E*.*crassipes* leaf extract (blue) Collagen type2 and (red) 51 Col-ECEE matrix

The type 2 collagen spectrum (blue) showed characteristic amide peaks: Amide A at ∼3325 cm^−1^ (N-H stretching of peptide linkages), Amide I at ∼1637 cm^−1^ (C=O stretching) and Amide II near 1550 cm^−1^ (N-H bending and C-N stretching). These are typical of the protein backbone of collagen. Minor peaks at ∼1450 cm^−1^ (C-H bending) and ∼1235 cm^−1^ (Amide III) were also present, supporting the structural integrity of collagen. The FTIR spectrum of the 5:1 Col-ECEE matrix (red) showed significant changes: the broad O-H/N-H stretching peak remained at 3325.28 cm^−1^, confirming hydroxyl and amide group presence post-conjugation. A shift of the amide I peak from 1656.85 cm^−1^ in the extract to 1637.56 cm^−1^ in the matrix suggested interactions between the extract’s carbonyl/hydroxyl groups and collagen’s amide bonds. The appearance or enhancement of peaks at 1439.97 cm^−1^ (C-H bending), 1010.70 cm^−1^ (C-O stretching), 950.91 cm^−1^ and 555.50 cm^−1^ indicated new bonding interactions, possibly hydrogen bonding or covalent modifications.These spectral changes confirm successful conjugation between collagen and the phytochemicals present *in E. crassipes* extract. Particularly, the shift and intensity differences in the amide I and O-H regions strongly suggest hydrogen bonding or electrostatic interactions, enhancing matrix stability. The alterations in the <1000 cm^−1^ fingerprint region also reflect new structural arrangements due to conjugation. Thus, FTIR analysis validates that functional groups such as hydroxyl, carbonyl and amine groups are involved in forming the Col-ECEE matrix, supporting its potential for biomedical applications like wound healing due to improved stability and bioactivity.

The scanning electron microscopy (SEM) analysis (Fig. 10) of the 5:1 Col-ECEE matrix revealed the formation of a sheet-like structure, indicative of a well-formed matrix. This morphology suggested that the *E. crassipes* leaf extract had effectively dispersed within the collagen matrix, forming continuous and cohesive layers. The porous layered surface of the matrix enhances the drug loading capacity and potentially improves the drug interaction with biological targets. The presence of a well-defined sheet structure implied that the matrix was mechanically stable, which was essential for biomedical applic ations, such as drug delivery systems or wound healing materials, where structural integrity was crucial. The sheet-like stable Col-ECEE matrix was successfully developed with enhanced functionality in therapeutic applications.

**Figure 10.**
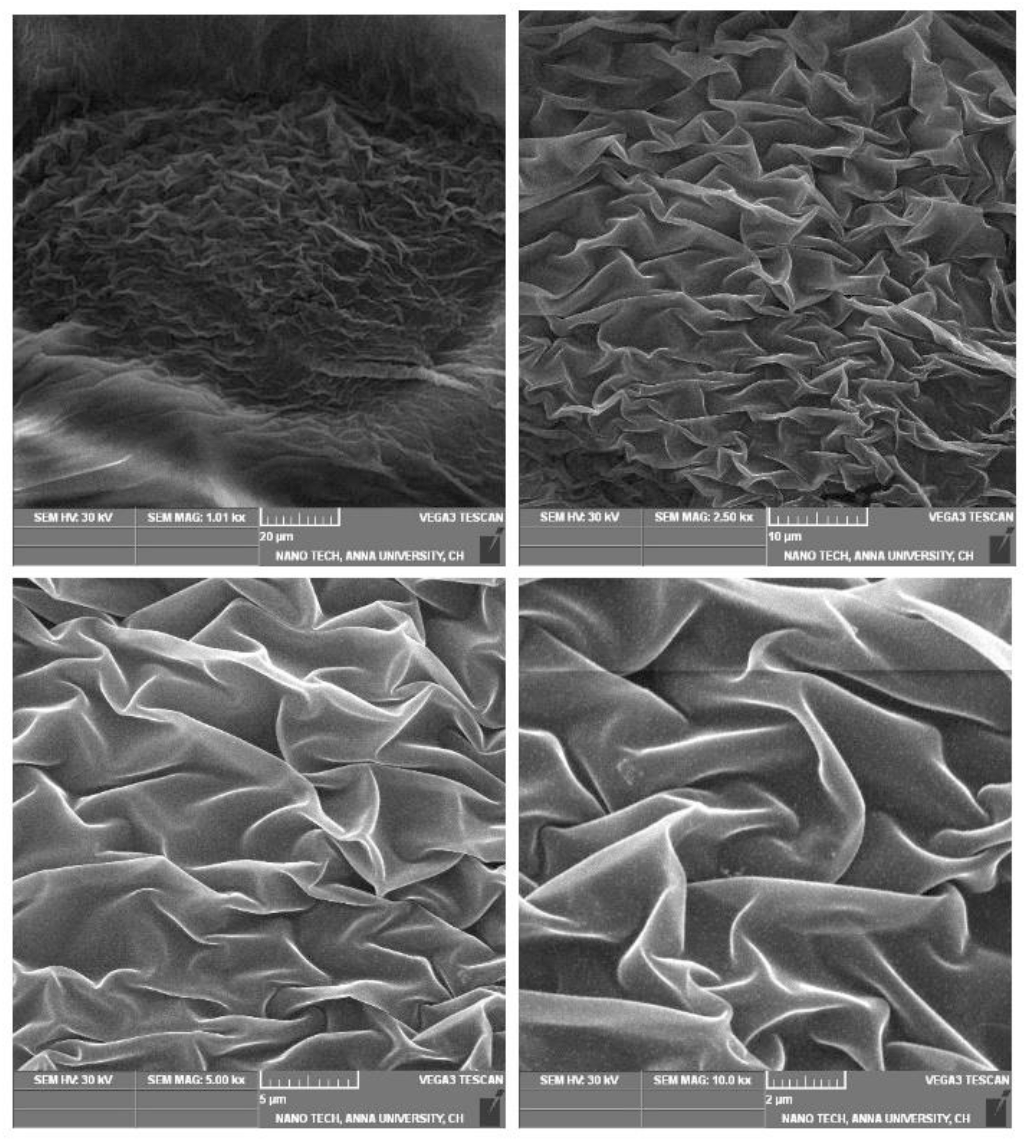
SEM image of 51 Col-ECEE matrix

The GC-MS analysis of ethanol extract of *E. crassipes* leaf identified ten bioactive compounds, listed in Table 1 with their respective IUPAC names and retention times and the structures are depicted in Fig. 11. These bioactive compounds were further analyzed through docking studies, where scoring functions and the number of hydrogen bonds formed with surrounding amino acids were used to predict their binding modes, binding affinities and molecular orientations.

**Table 1.**
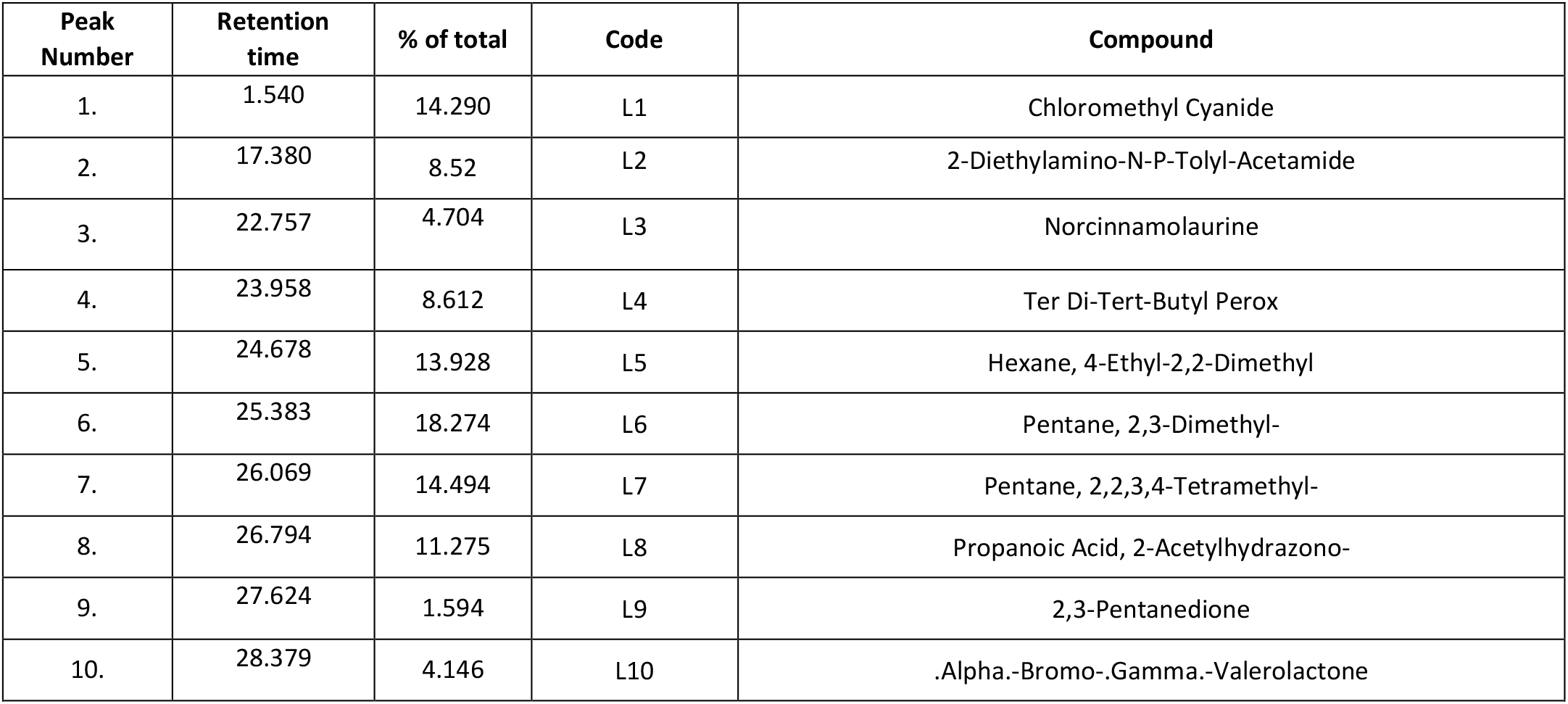
GC-MS analysis of ethanolic extract of *E. crassipes* leaf.

| Peak Number | Retention time | % of total | Code | Compound |
| --- | --- | --- | --- | --- |
| 1. | 1.540 | 14.290 | L1 | Chloromethyl Cyanide |
| 2. | 17.380 | 8.52 | L2 | 2-Diethylamino-N-P-Tolyl-Acetamide |
| 3. | 22.757 | 4.704 | L3 | Norcinnamolaurine |
| 4. | 23.958 | 8.612 | L4 | Ter Di-Tert-Butyl Perox |
| 5. | 24.678 | 13.928 | L5 | Hexane, 4-Ethyl-2,2-Dimethyl |
| 6. | 25.383 | 18.274 | L6 | Pentane, 2,3-Dimethyl- |
| 7. | 26.069 | 14.494 | L7 | Pentane, 2,2,3,4-Tetramethyl- |
| 8. | 26.794 | 11.275 | L8 | Propanoic Acid, 2-Acetylhydrazono- |
| 9. | 27.624 | 1.594 | L9 | 2,3-Pentanedione |
| 10. | 28.379 | 4.146 | L10 | .Alpha.-Bromo-.Gamma.-Valerolactone |
| Table 1 GC-MS analysis of ethanolic extract of *E. crassipes* leaf |  |  |  |  |

**Figure 11.**
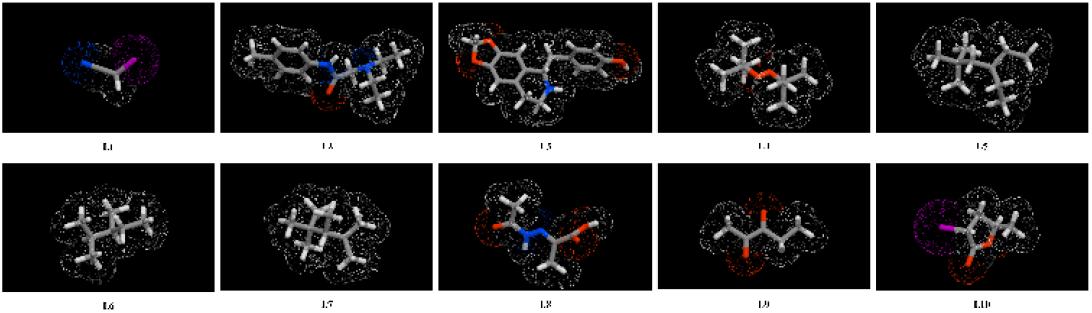
Structures of bioactive compounds of *E. crassipes* leaf extract

The binding energy scores for all the bioactive compounds identified in the *E. crassipes* leaf extract are presented in Table 2. A higher LibDock score signifies a stronger receptor-ligand binding affinity (Alam and Khan, 2018).

**Table 2.** Molecular docking binding energy score of bioactive compounds of *E. crassipes* leaf extract.

| Sl No | Compounds | LibDock score |  |  |  |
| --- | --- | --- | --- | --- | --- |
|  |  | 7WEU | 3BAJ | 2QLY | 3KBX |
| 1 | Chloromethyl Cyanide | 31.7551 | 31.2179 | 28.6104 | 34.3198 |
| 2 | 2-Diethylamino-N-P-Tolyl-Acetamide | 92.7455 | 81.4528 | 102.539 | 100.74 |
| 3 | Norcinnamolaurine<br>(Norcinnamolaurine) | 90.8323 | 105.335 | 102.766 | 116.732 |
| 4 | Ter Di-Tert-Butyl Perox | 56.2347 | 45.6842 | 55.5327 | 64.8152 |
| 5 | Hexane, 4-Ethyl-2,2-Dimethyl | 58.9244 | 52.9154 | 63.6255 | 66.8728 |
| 6 | Pentane, 2,3-Dimethyl- | 46.7843 | 43.5312 | 48.5603 | 51.5288 |
| 7 | Pentane, 2,2,3,4-Tetramethyl- | 55.7196 | 42.4137 | 48.2335 | 58.0587 |
| 8 | Propanoic Acid, 2-Acetylhydrazono- | 73.4009 | 61.4141 | 58.7313 | 75.9699 |
| 9 | 2,3-Pentanedione | 50.183 | 48.3229 | 43.3639 | 54.9281 |
| 10 | .Alpha.-Bromo-.Gamma.-Valerolactone | 53.4266 | 48.8677 | 50.1613 | 58.9277 |

For the target protein ‘7WEU,’ 2-Diethylamino-N-P-Tolyl-Acetamide achieved a LibDock score of 92.7455, indicating notable binding affinity, with three conventional hydrogen bonds formed (Table 3). Norcinnamolaurine exhibited the highest LibDock scores of 105.335, 102.766 and 116.732 for the targets ‘3BAJ’, ‘2QLY’ and ‘3KBX’, respectively, forming two conventional hydrogen bonds with each target (Table 3). The interactions and amino acids binding sites between the targets, ‘7WEU’, ‘3BAJ’, ‘2QLY’ and ‘3KBX’ and the selected ligands, 2-Diethylamino-N-P-Tolyl-Acetamide and Norcinnamolaurine were illustrated in Fig. 12-15. The receptor surface interaction of the ligand scrutinized by Aromatic, Interpolated Charge, H-Bonds, Hydrophobicity, Ionizability, SAS (Solvent Accessibility Surface) parameters were also depicted in Fig. 12-15.

**Table 3.**
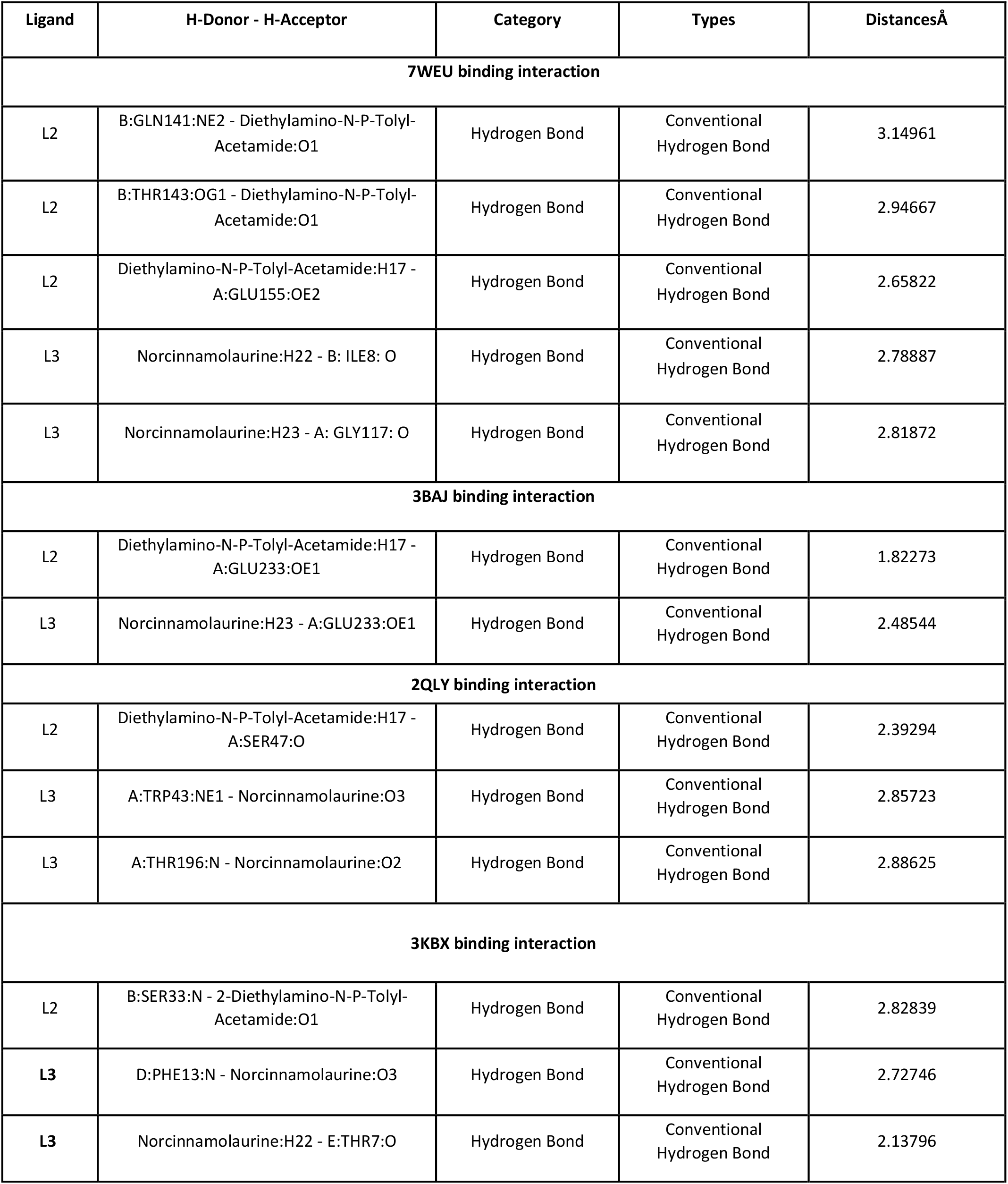
binding interaction.

| Ligand | H-Donor - H-Acceptor | Category | Types | DistancesÅ |
| --- | --- | --- | --- | --- |
| **7WEU binding interaction** |  |  |  |  |
| L2 | B:GLN141:NE2 - Diethylamino-N-P-Tolyl-Acetamide:O1 | Hydrogen Bond | Conventional Hydrogen Bond | 3.14961 |
| L2 | B:THR143:OG1 - Diethylamino-N-P-Tolyl-Acetamide:O1 | Hydrogen Bond | Conventional Hydrogen Bond | 2.94667 |
| L2 | Diethylamino-N-P-Tolyl-Acetamide:H17 - A:GLU155:OE2 | Hydrogen Bond | Conventional Hydrogen Bond | 2.65822 |
| L3 | Norcinnamolaurine:H22 - B: ILE8: O | Hydrogen Bond | Conventional Hydrogen Bond | 2.78887 |
| L3 | Norcinnamolaurine:H23 - A: GLY117: O | Hydrogen Bond | Conventional Hydrogen Bond | 2.81872 |
| **3BAJ binding interaction** |  |  |  |  |
| L2 | Diethylamino-N-P-Tolyl-Acetamide:H17 - A:GLU233:OE1 | Hydrogen Bond | Conventional Hydrogen Bond | 1.82273 |
| L3 | Norcinnamolaurine:H23 - A:GLU233:OE1 | Hydrogen Bond | Conventional Hydrogen Bond | 2.48544 |
| **2QLY binding interaction** |  |  |  |  |
| L2 | Diethylamino-N-P-Tolyl-Acetamide:H17 - A:SER47:O | Hydrogen Bond | Conventional Hydrogen Bond | 2.39294 |
| L3 | A:TRP43:NE1 - Norcinnamolaurine:O3 | Hydrogen Bond | Conventional Hydrogen Bond | 2.85723 |
| L3 | A:THR196:N - Norcinnamolaurine:O2 | Hydrogen Bond | Conventional Hydrogen Bond | 2.88625 |
| **3KBX binding interaction** |  |  |  |  |
| L2 | B:SER33:N - 2-Diethylamino-N-P-Tolyl-Acetamide:O1 | Hydrogen Bond | Conventional Hydrogen Bond | 2.82839 |
| L3 | D:PHE13:N - Norcinnamolaurine:O3 | Hydrogen Bond | Conventional Hydrogen Bond | 2.72746 |
| L3 | Norcinnamolaurine:H22 - E:THR7:O | Hydrogen Bond | Conventional Hydrogen Bond | 2.13796 |
| **Table 3 binding interaction** |  |  |  |  |

**Figure 12.**
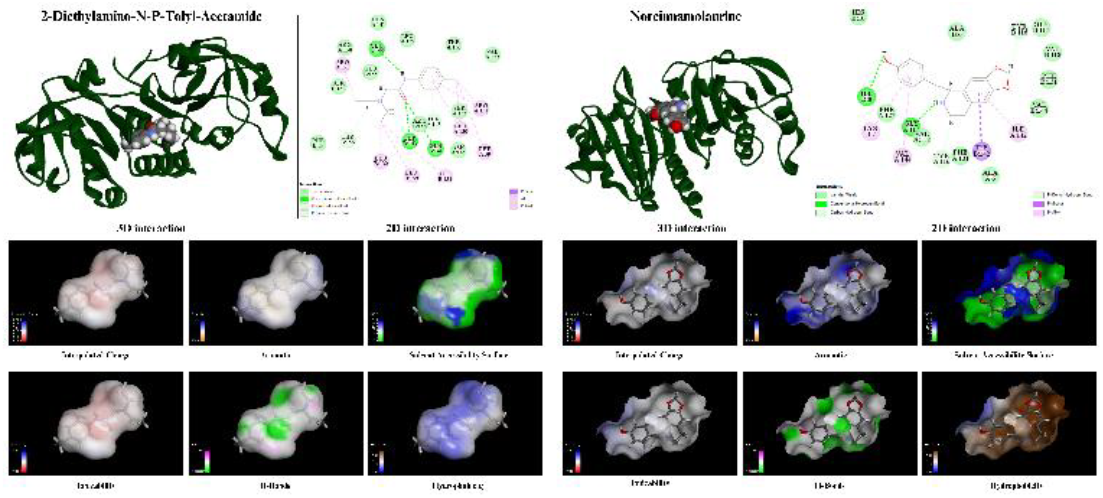
3D and 2D Molecular docking binding interaction of 2-Diethylamino-N-P-Tolyl-Acetamide (LEFT) and Norcinnamolaurine (RIGHT) on 7WEU and their surface interaction on 7WEU

**Figure 13.**
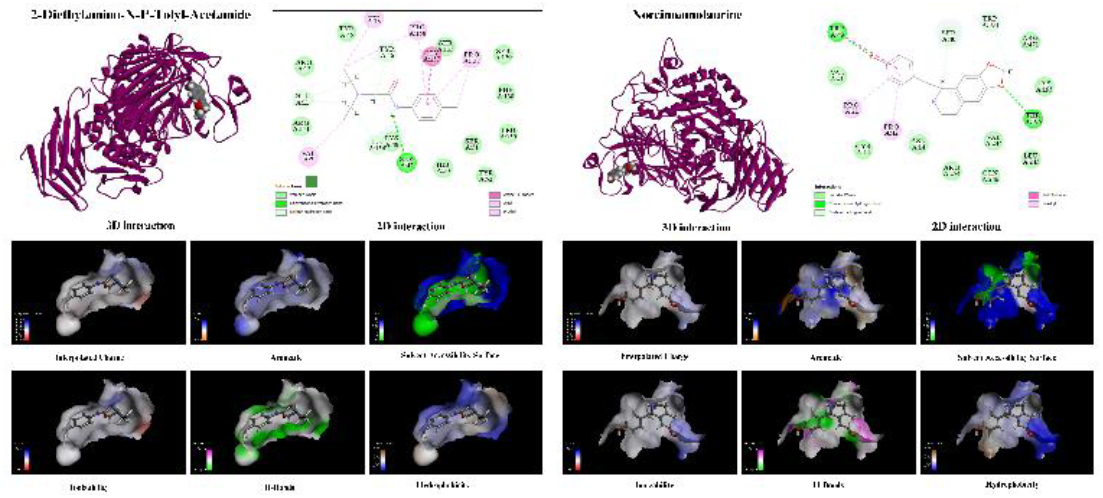
3D and 2D Molecular docking binding interaction of 2-Diethylamino-N-P-Tolyl-Acetamide (LEFT) and Norcinnamolaurine (RIGHT) on 2QLY and their surface interaction on 2QLY

**Figure 14.**
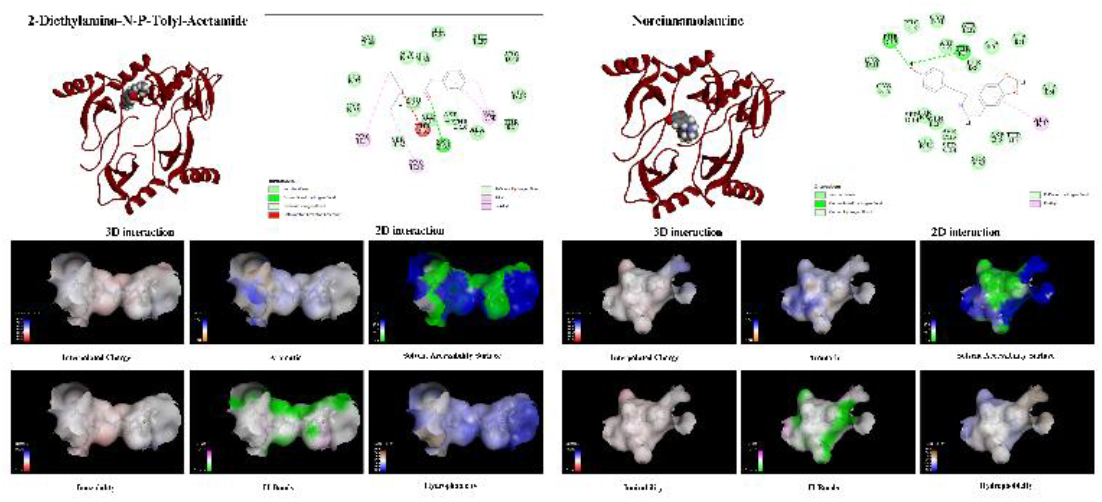
3D and 2D Molecular docking binding interaction of 2-Diethylamino-N-P-Tolyl-Acetamide (LEFT) and Norcinnamolaurine (RIGHT) on 3KBX and their surface interaction on 3KBX

**Figure 15.**
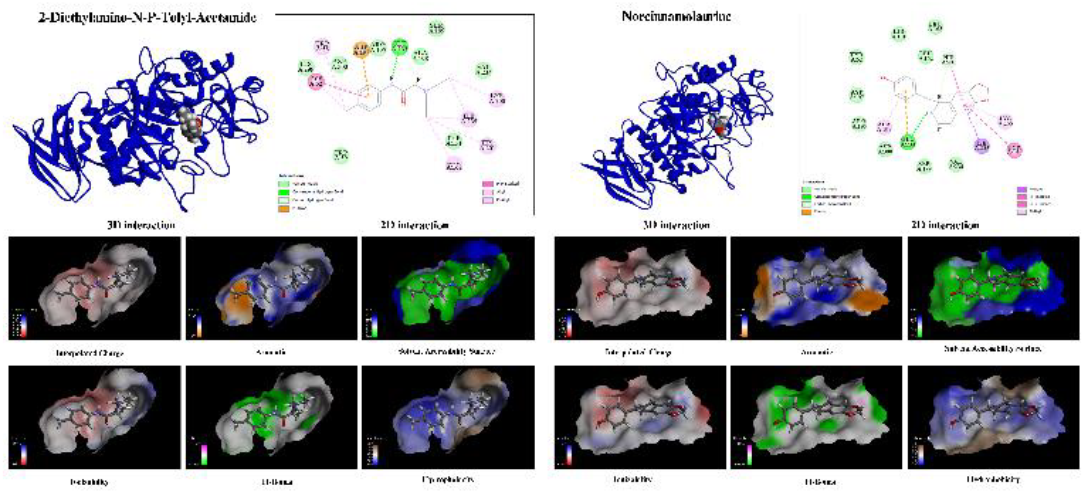
3D and 2D Molecular docking binding interaction of 2-Diethylamino-N-P-Tolyl-Acetamide (LEFT) and Norcinnamolaurine (RIGHT) on 3BAJ and their surface interaction on 3BAJ

AlogP98 and 2D polar surface area served as key descriptors in developing the ADMET model. This study includes the 99% and 95% confidence limit ellipses, which outline the regions indicating suitable intestinal absorption and blood-brain barrier (BBB) penetration. In Figure 16, the 99% confidence limits for intestinal absorption and BBB are marked in green and blue, while the 95% confidence limits are shown in red and pink, respectively. According to the ADMET model results, eight compounds fell within the 95% confidence limits for intestinal absorption, with two compounds lying outside this boundary (Fig. 16).

**Figure 16.**
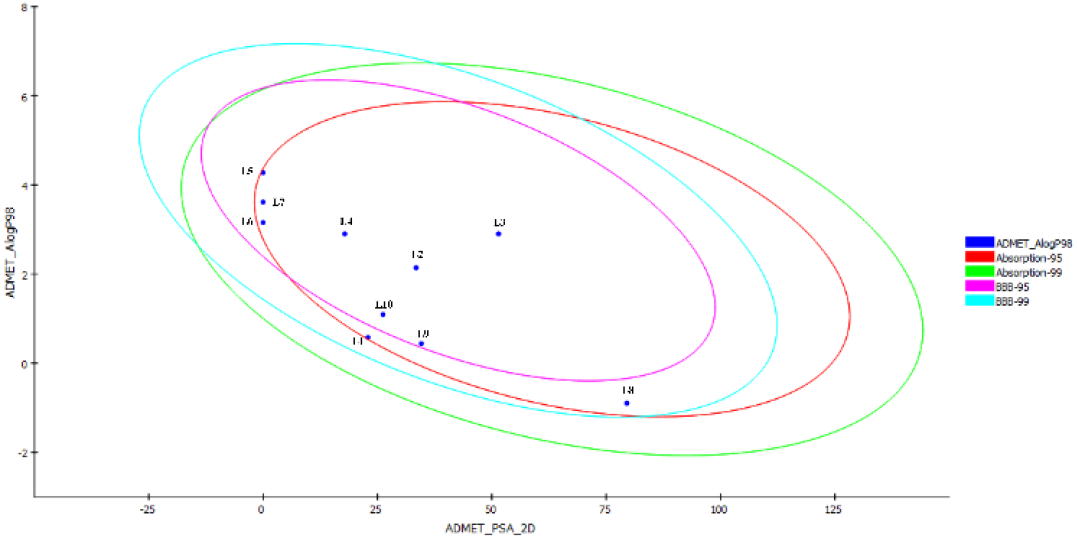
2D polar surface area of bioactive compounds of *E. crassipes* leaf extract

Based on 3D and 2D molecular docking interaction studies, the compounds 2-Diethylamino-N-P-Tolyl-Acetamide and Norcinnamolaurine, identified from the GC-MS profile of *Eichhornia crassipes* leaf extract, were selected as lead bioactive candidates for wound healing evaluation. Their strong binding affinity toward wound-related target proteins supports the potential therapeutic relevance of the extract.

The ADME analysis of 2-Diethylamino-N-P-Tolyl-Acetamide and Norcinnamolaurine revealed their absorption, distribution, metabolism and excretion characteristics (Table 4). Both compounds possessed physicochemical properties with over two rotatable bonds and contained hydrogen bond donor and acceptor sites. Their water solubility, based on Log S (ESOL) values, indicated that both compounds are soluble. In terms of pharmacokinetics, Norcinnamolaurine acts as an inhibitor of CYP1A2 (linked to obsessive-compulsive disorder), CYP2D6 (an enzyme influencing drug metabolism) and CYP3A4 (an enzyme involved in the oxidation of small foreign organic molecules like drugs and toxins). Similarly, 2-Diethylamino-N-P-Tolyl-Acetamide serves as a CYP2D6 inhibitor. Regarding drug-likeness, both compounds adhered to Lipinski’s rule of five, with no violations and each has a bioavailability score of 0.55. This score suggests that, based on the rule, at least 10% of each compound would likely be bioavailable in rats.

**Table 4.**
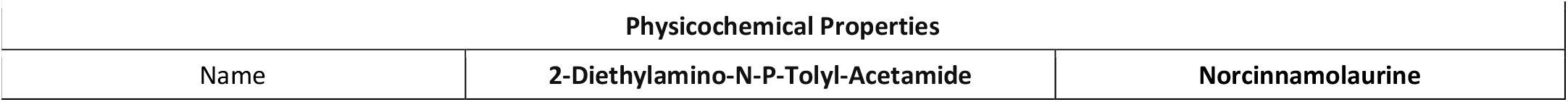
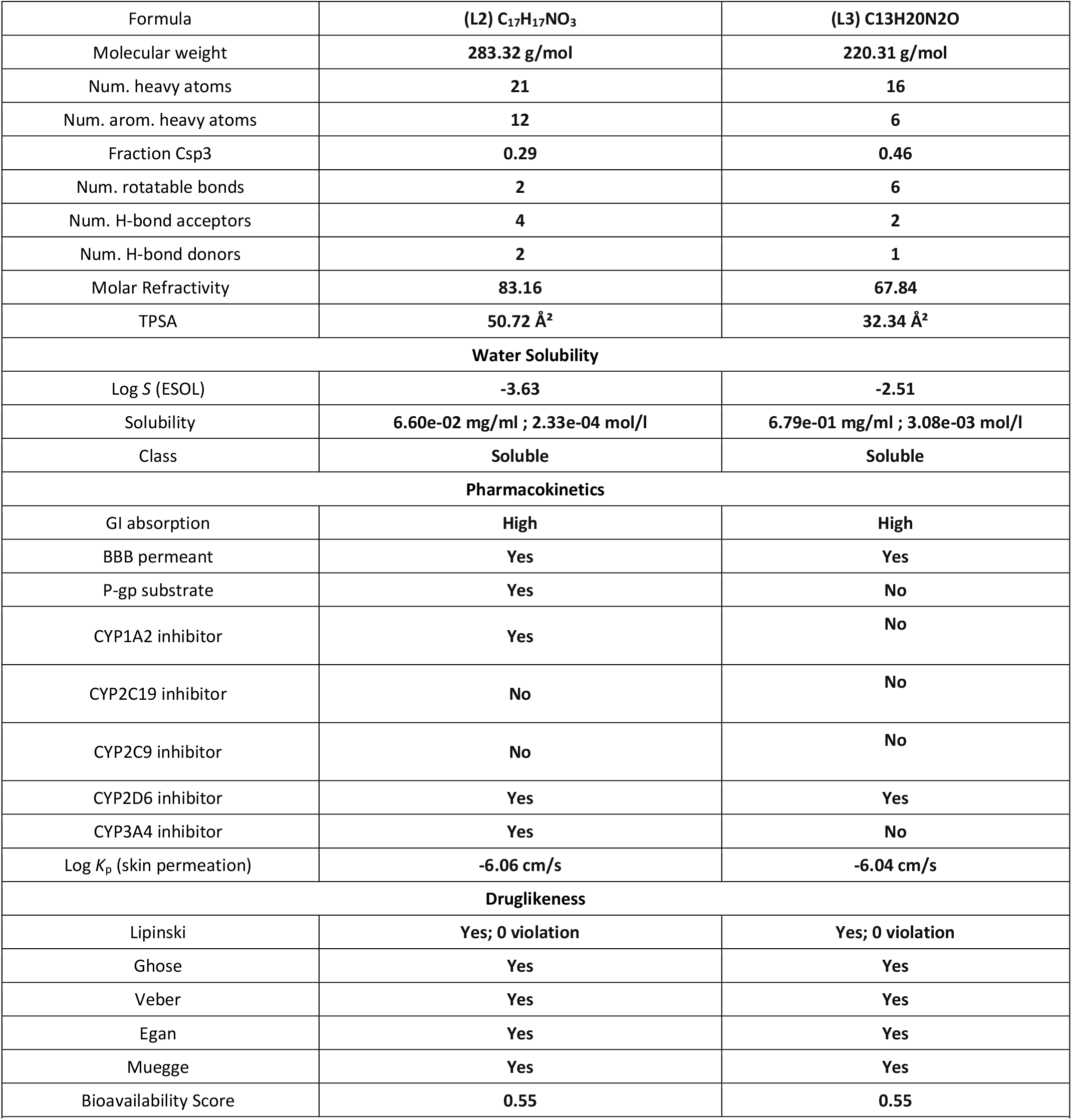
ADME analysis of 2-Diethylamino-N-P-Tolyl-Acetamide and Norcinnamolaurine.

The TOPKAT (Toxicity Prediction by Komputer-Assisted Technology) method was applied to assess the therapeutic compatibility and predict toxicity of the drug using *In silico* models (Table 5). TOPKAT is a valuable tool for quantitative toxicity predictions, utilized in quantitative structure-activity relationship (QSAR) models. Through these QSAR models, TOPKAT calculated the probability values to evaluate toxicity and adhered to the optimal predictive space (OPS) criterion. Results outside OPS are considered unreliable, potentially indicating false positives. The Ames probability values classify toxicity levels: 0.0 to 0.30 (non-toxic), 0.30 to 0.70 (intermediate) and 0.70 to 1.0 (toxic) (Tanwar et al., 2019). Additional descriptors include Ames mutagenicity, enrichment and scores for reliability assessment. TOPKAT also provides values for carcinogenic potency (TD50), rat oral LD50, rat inhalational LC50 and Daphnia EC50. Higher TD50, LD50, LC50 and EC50 values indicates reduced toxicity and a higher safety index, enhancing the therapeutic potential of drug.

**Table 5.**
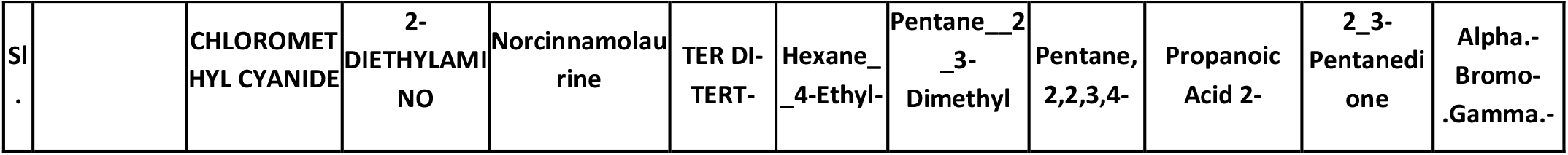
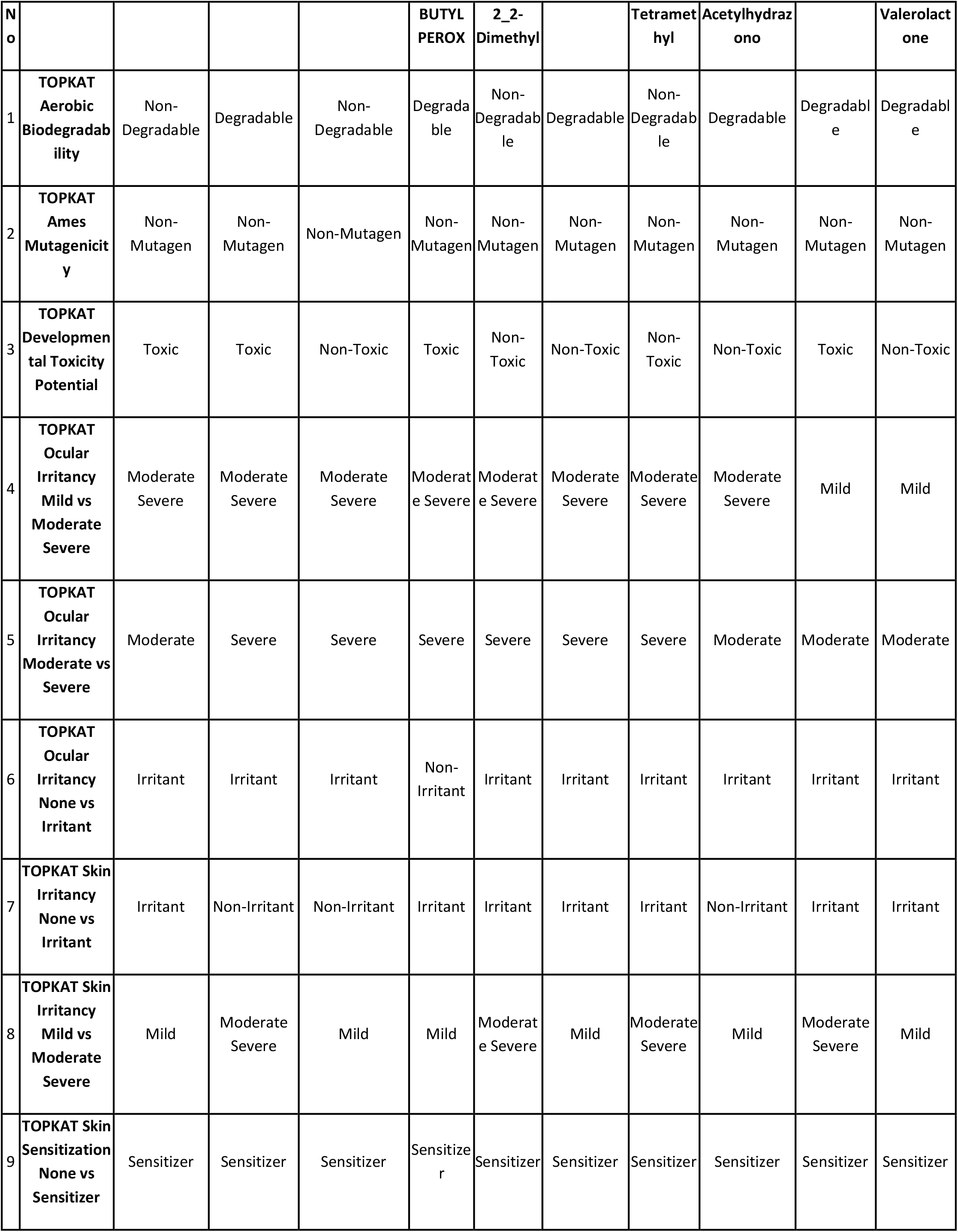
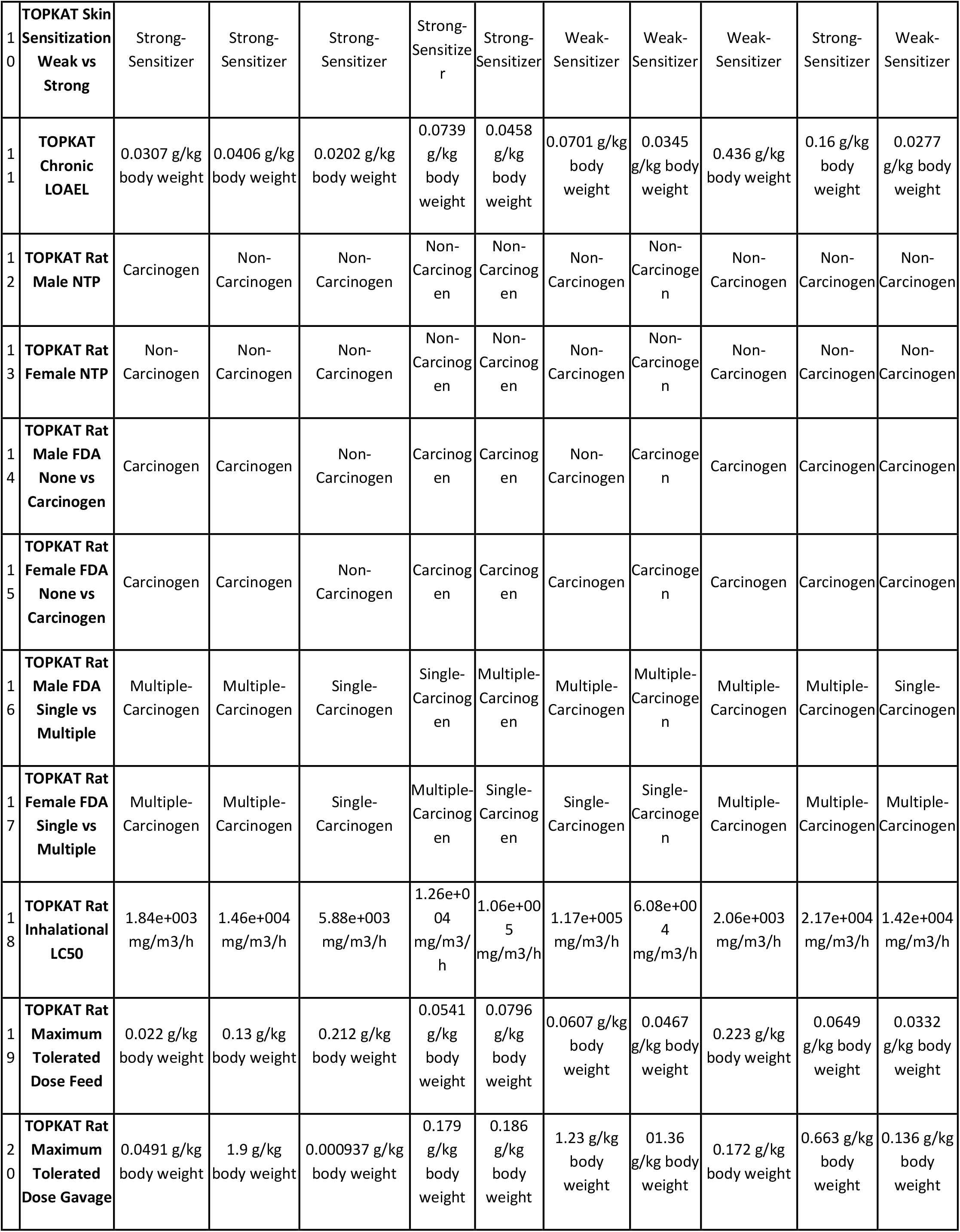
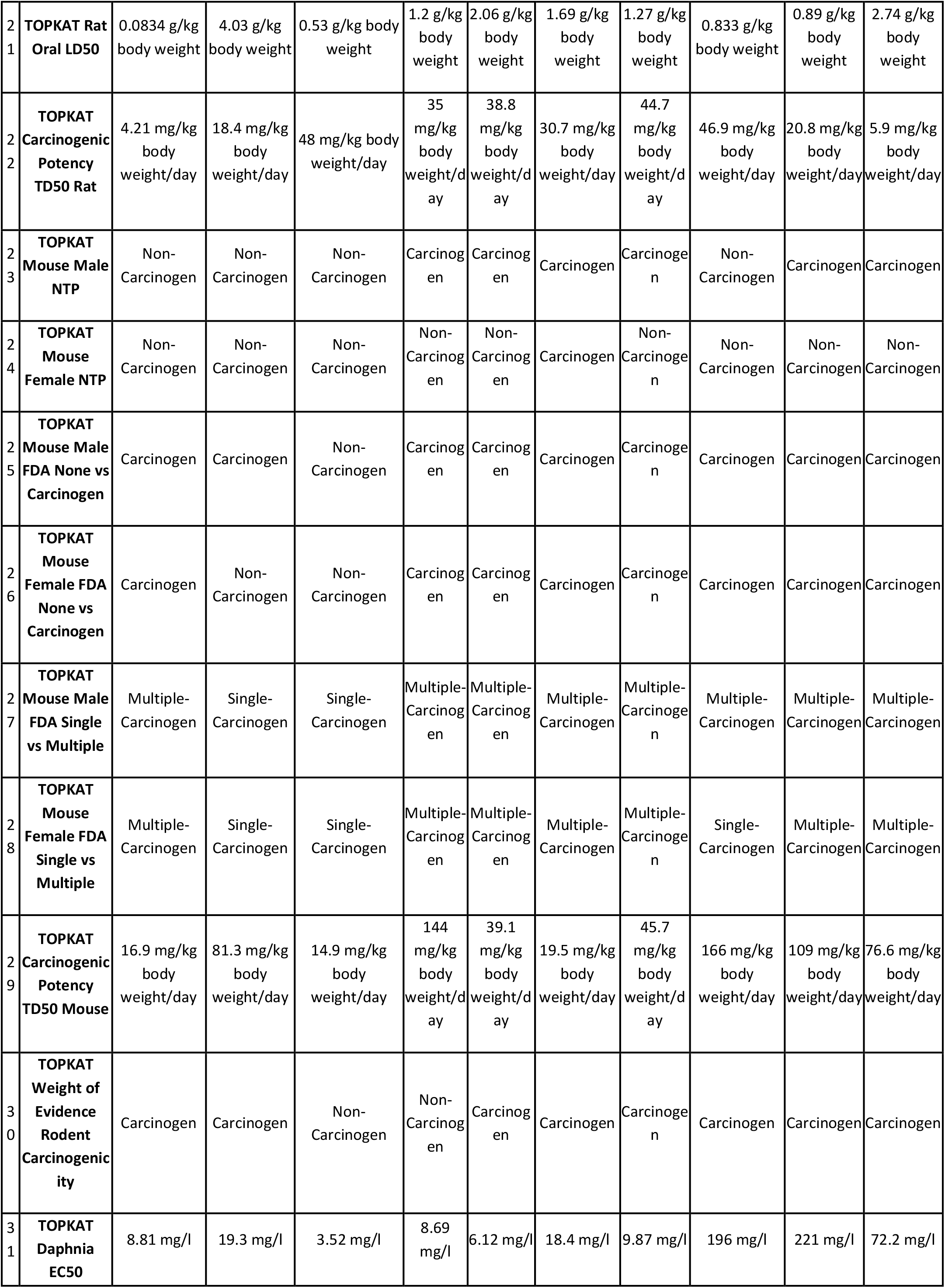
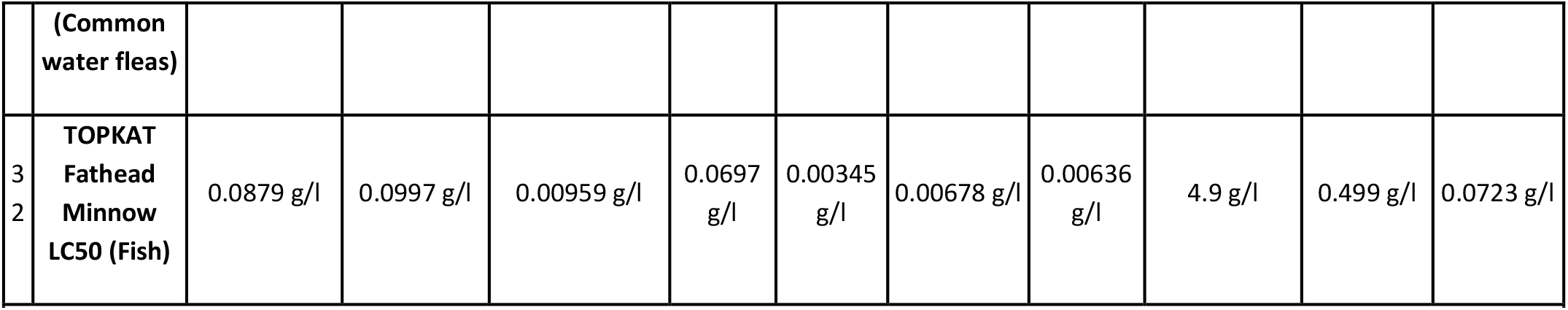
TOPKAT analysis of bioactive compounds of *E. crassipes* leaf extract.

Drug-like properties of the compounds identified in *E. crassipes* were evaluated based on Lipinski’s rule of five using the Molinspiration cheminformatic tool (Raju et al., 2022). As shown in Table 6, none of the compounds violated Lipinski’s rule, which states that a molecule was considered drug-like if it meets at least two of the following criteria: lipophilicity (Log P) below 5, molecular weight under 500 Daltons, fewer than 5 hydrogen donors and so on (Lipinski, 2004). It was noted that for natural products, lipophilicity may not always align with physicochemical profiles, occasionally deviating from the rule of five (Lipinski, 2016). The predicted Topological Polar Surface Area (TPSA) values of all compounds ranged between 18.47–78.76 Å^2^, indicating favourable oral bioavailability (Reena Roy et al., 2020). Compound flexibility, as shown by the number of rotatable bonds (ranging from 0 to 5), also correlates with bioavailability, which was assessed to further understand drug-like properties (Lipin et al., 2021).

**Table 6.**
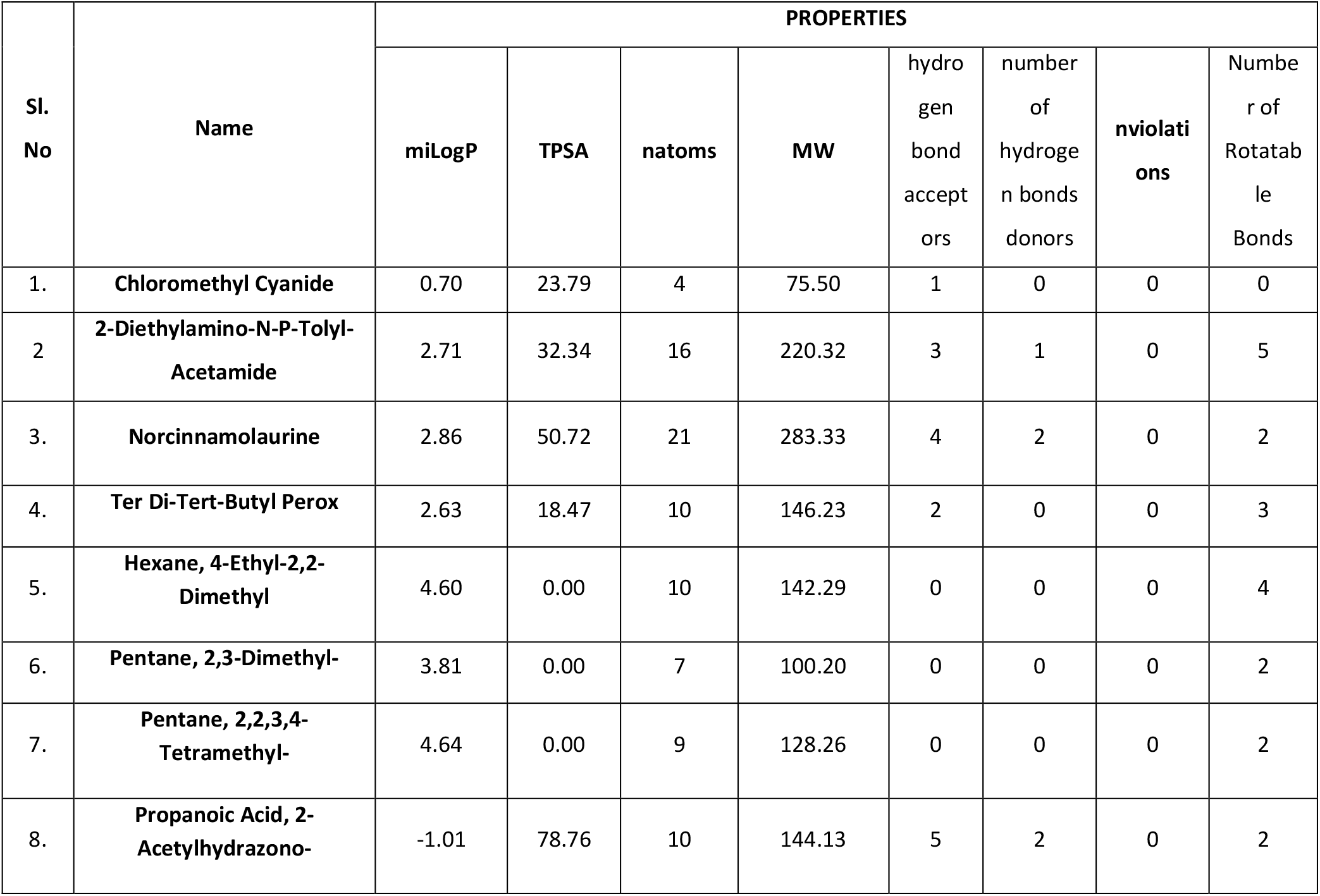
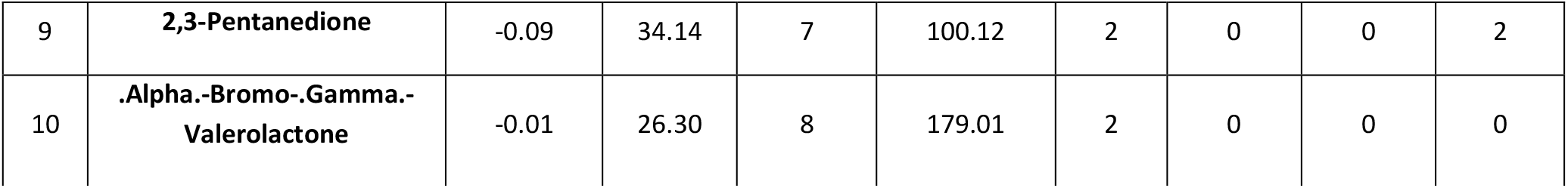
Drug-likeness scores of bioactive compounds of *E. crassipes* leaf extract.

The bioactivity scores of the *E. crassipes* compounds obtained from Molinspiration indicated that scores above 0.00 suggest a high level of biological activity, while scores between 0.5 and −0.5 reflect moderate activity and scores below −0.5 indicate inactivity (Reena Roy et al., 2020). As shown in Table 7, Norcinnamolaurine demonstrated a high level of bioactivity as a GPCR ligand, ion channel modulator and enzyme inhibitor, with scores of 0.25, 0.12 and 0.10, respectively. In contrast, 2-Diethylamino-N-P-Tolyl-Acetamide exhibited moderate activity, while the remaining compounds displayed inactivity in at least one bioactivity domain.

**Table 7.**
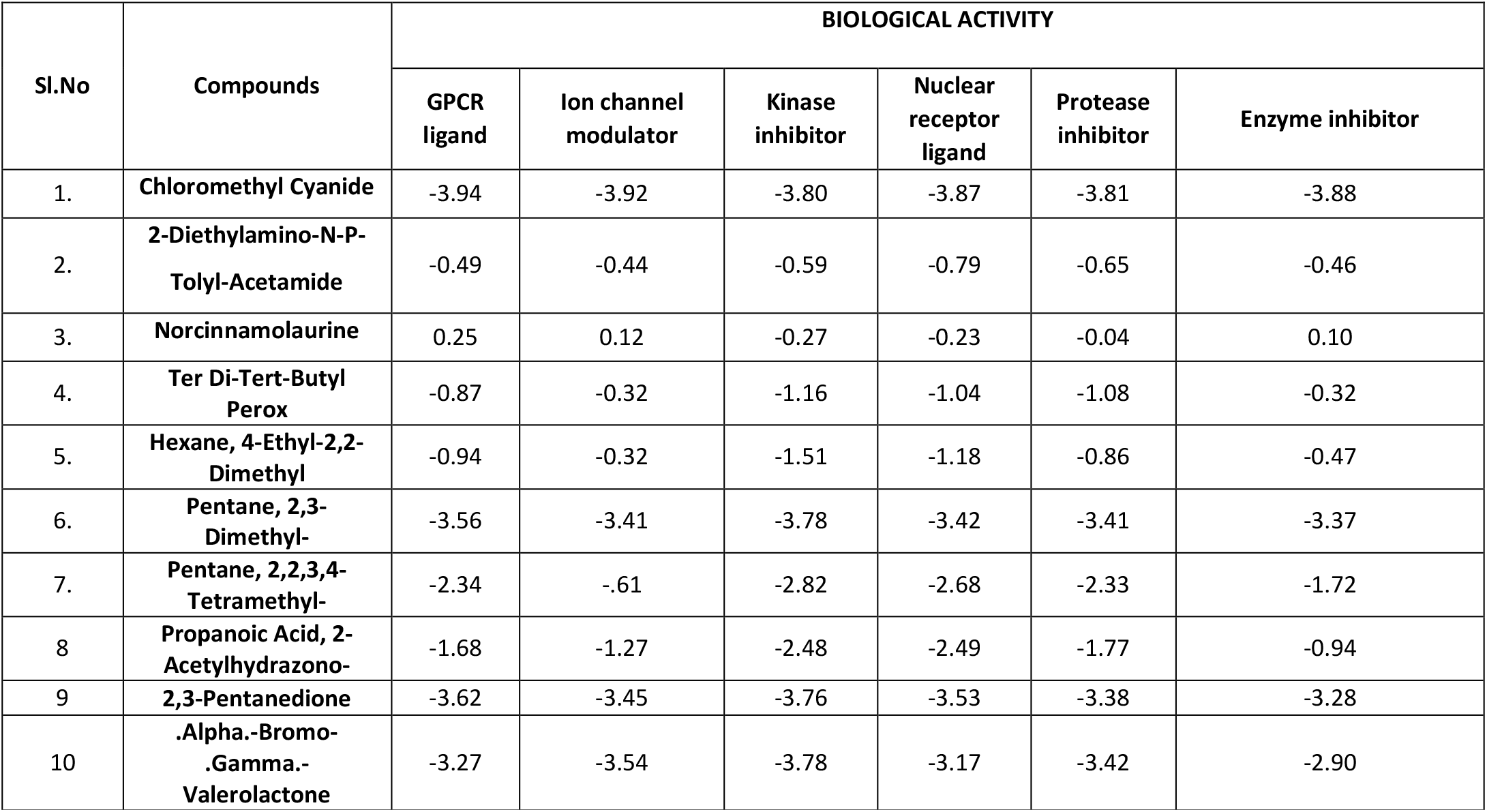
Biological activity scores of bioactive compounds of *E. crassipes* leaf extract.

The variation in extract concentrations across experiments was intentional, based on the biological context of each assay. Antioxidant and enzyme inhibition assays required broader ranges to determine IC_50_ values, while cytotoxicity and wound healing assays used lower or fixed concentrations to simulate physiologically relevant exposures and to avoid overt toxicity. This strategic variation enabled optimal data generation for each test system.

## Conclusion

The pursuit of high target potency requires a strategic approach that considers its relevance to efficacy and efficiency. This study introduces a novel bioactive wound healing matrix composed of collagen type 2 integrated with phytochemicals derived from *Eichhornia crassipes*, an underutilized and invasive aquatic species, as wound-healing agents and demonstrated the potential to develop innovative and versatile drugs. The research was distinctive in its multidisciplinary approach, combining phytochemical profiling (via GC-MS), molecular modeling (LibDock Score, ADMET and TOPKAT etc.) and comprehensive *in vitro* pharmacological evaluations, including antioxidant, antidiabetic, anti-inflammatory, antibacterial, cytotoxicity and wound closure assays. The *In vitro* analysis of *E. crassipes* showed promising antioxidant, antidiabetic and anti-inflammatory activity, highlighting its potential across multiple fields. The Col-ECEE matrix showed enhanced cell viability and significant *In vitro* wound healing potential, establishing it as a strong candidate for therapeutic wound care applications. The successful conjugation of the extract to collagen, confirmed via FTIR and SEM, enhances the structural stability and functional performance of the matrix. This work pioneers the transformation of *E. crassipes* into a pharmacologically active and biocompatible wound dressing, offering an environmentally sustainable and scientifically validated approach for biomaterial-based therapeutics. Future research, including specific mechanisms, *in vivo* testing and clinical trials, would be essential to fully validate its effectiveness and suitability in practical wound healing treatments. By identifying and characterizing key phytochemical descriptors responsible for cytotoxic activity and *In vitro* wound healing through QSAR modelling, a deeper understanding of their functional roles was gained. Top compounds from *E. crassipes* were screened *in silico* and their efficacy was verified through oral bioavailability assessment, docking, ADMET risk screening and *in silico* pharmacokinetics/pharmacodynamics (PK/PD) analysis. Systems pharmacology and structure-guided insights into molecular interactions provided an understanding of on- and off-target effects and the cell networks involved. Notably, two GC-MS–identified compounds—2-Diethylamino-N-P-Tolyl-Acetamide and Norcinnamolaurine were evaluated for their potential anti-inflammatory, antioxidant and antidiabetic activities. Their strong binding affinities with multiple wound-associated targets suggest that these compounds may significantly contribute to the observed pharmacological effects of the *E. crassipes* extract. These compounds (Norcinnamolaurine and 2-Diethylamino-N-P-Tolyl-Acetamide) emerged as potential therapeutic molecules, with docking studies suggesting strong molecular interactions and signaling pathway involvement. This *in silico* approach paves the way for a streamlined drug discovery process, potentially minimizing the need for extensive *In vitro* and *in vivo* studies due to comprehensive PK/PD patterning, which aids in efficient drug screening.

## Author contributions

S.VENKATA KRISHNAN – Conceptualization, Data curation, Formal analysis, Funding acquisition, Investigation, Methodology, Project administration, Software, Visualization, Writing – original draft, Writing – review & editing.

## Data availability

The authors confirm that the data supporting the findings of this study are available within the article which has been included as part of Figures or Graphs and Table

---

**Table 1.** GC-MS analysis of ethanolic extract of *E. crassipes* leaf.

| Peak Number | Retention time | % of total | Code | Compound |
| --- | --- | --- | --- | --- |
| 1. | 1.540 | 14.290 | L1 | Chloromethyl Cyanide |
| 2. | 17.380 | 8.52 | L2 | 2-Diethylamino-N-P-Tolyl-Acetamide |
| 3. | 22.757 | 4.704 | L3 | Norcinnamolaurine |
| 4. | 23.958 | 8.612 | L4 | Ter Di-Tert-Butyl Perox |
| 5. | 24.678 | 13.928 | L5 | Hexane, 4-Ethyl-2,2-Dimethyl |
| 6. | 25.383 | 18.274 | L6 | Pentane, 2,3-Dimethyl- |
| 7. | 26.069 | 14.494 | L7 | Pentane, 2,2,3,4-Tetramethyl- |
| 8. | 26.794 | 11.275 | L8 | Propanoic Acid, 2-Acetylhydrazono- |
| 9. | 27.624 | 1.594 | L9 | 2,3-Pentanedione |
| 10. | 28.379 | 4.146 | L10 | .Alpha.-Bromo-.Gamma.-Valerolactone |

**Table 2.** Molecular docking binding energy score of bioactive compounds of *E. crassipes* leaf extract.

| Sl No | Compounds | LibDock score |  |  |  |
| --- | --- | --- | --- | --- | --- |
|  |  | 7WEU | 3BAJ | 2QLY | 3KBX |
| 1 | Chloromethyl Cyanide | 31.7551 | 31.2179 | 28.6104 | 34.3198 |
| 2 | 2-Diethylamino-N-P-Tolyl-Acetamide | 92.7455 | 81.4528 | 102.539 | 100.74 |
| 3 | Norcinnamolaurine<br>(Norcinnamolaurine) | 90.8323 | 105.335 | 102.766 | 116.732 |
| 4 | Ter Di-Tert-Butyl Perox | 56.2347 | 45.6842 | 55.5327 | 64.8152 |
| 5 | Hexane, 4-Ethyl-2,2-Dimethyl | 58.9244 | 52.9154 | 63.6255 | 66.8728 |
| 6 | Pentane, 2,3-Dimethyl- | 46.7843 | 43.5312 | 48.5603 | 51.5288 |
| 7 | Pentane, 2,2,3,4-Tetramethyl- | 55.7196 | 42.4137 | 48.2335 | 58.0587 |
| 8 | Propanoic Acid, 2-Acetylhydrazono- | 73.4009 | 61.4141 | 58.7313 | 75.9699 |
| 9 | 2,3-Pentanedione | 50.183 | 48.3229 | 43.3639 | 54.9281 |
| 10 | .Alpha.-Bromo-.Gamma.- | 53.4266 | 48.8677 | 50.1613 | 58.9277 |
|  | Valerolactone |  |  |  |  |

**Table 3.** binding interaction.

| Ligand | H-Donor - H-Acceptor | Category | Types | DistancesÅ |
| --- | --- | --- | --- | --- |
| **7WEU binding interaction** |  |  |  |  |
| L2 | B:GLN141:NE2 -<br>Diethylamino-N-P-Tolyl-Acetamide:O1 | Hydrogen Bond | Conventional<br>Hydrogen Bond | 3.14961 |
| L2 | B:THR143:OG1 -<br>Diethylamino-N-P-Tolyl-Acetamide:O1 | Hydrogen Bond | Conventional<br>Hydrogen Bond | 2.94667 |
| L2 | Diethylamino-N-P-Tolyl-Acetamide:H17 -<br>A:GLU155:OE2 | Hydrogen Bond | Conventional<br>Hydrogen Bond | 2.65822 |
| L3 | Norcinnamolaurine:H22 - B:<br>ILE8: O | Hydrogen Bond | Conventional<br>Hydrogen Bond | 2.78887 |
| L3 | Norcinnamolaurine:H23 - A:<br>GLY117: O | Hydrogen Bond | Conventional<br>Hydrogen Bond | 2.81872 |
| **3BAJ binding interaction** |  |  |  |  |
| L2 | Diethylamino-N-P-Tolyl-Acetamide:H17 -<br>A:GLU233:OE1 | Hydrogen Bond | Conventional<br>Hydrogen Bond | 1.82273 |
| L3 | Norcinnamolaurine:H23 -<br>A:GLU233:OE1 | Hydrogen Bond | Conventional<br>Hydrogen Bond | 2.48544 |
| **2QLY binding interaction** |  |  |  |  |
| L2 | Diethylamino-N-P-Tolyl-Acetamide:H17 - A:SER47:O | Hydrogen Bond | Conventional<br>Hydrogen Bond | 2.39294 |
| L3 | A:TRP43:NE1 -<br>Norcinnamolaurine:O3 | Hydrogen Bond | Conventional<br>Hydrogen Bond | 2.85723 |
| L3 | A:THR196:N -<br>Norcinnamolaurine:O2 | Hydrogen Bond | Conventional<br>Hydrogen Bond | 2.88625 |

| 3 KBX binding interaction |  |  |  |  |
| --- | --- | --- | --- | --- |
| L2 | B:SER33:N - 2-Diethylamino-N-P-Tolyl-Acetamide:O1 | Hydrogen Bond | Conventional Hydrogen Bond | 2.82839 |
| L3 | D:PHE13:N - Norcinnamolaurine:O3 | Hydrogen Bond | Conventional Hydrogen Bond | 2.72746 |
| L3 | Norcinnamolaurine:H22 - E:THR7:O | Hydrogen Bond | Conventional Hydrogen Bond | 2.13796 |

**Table 4.**
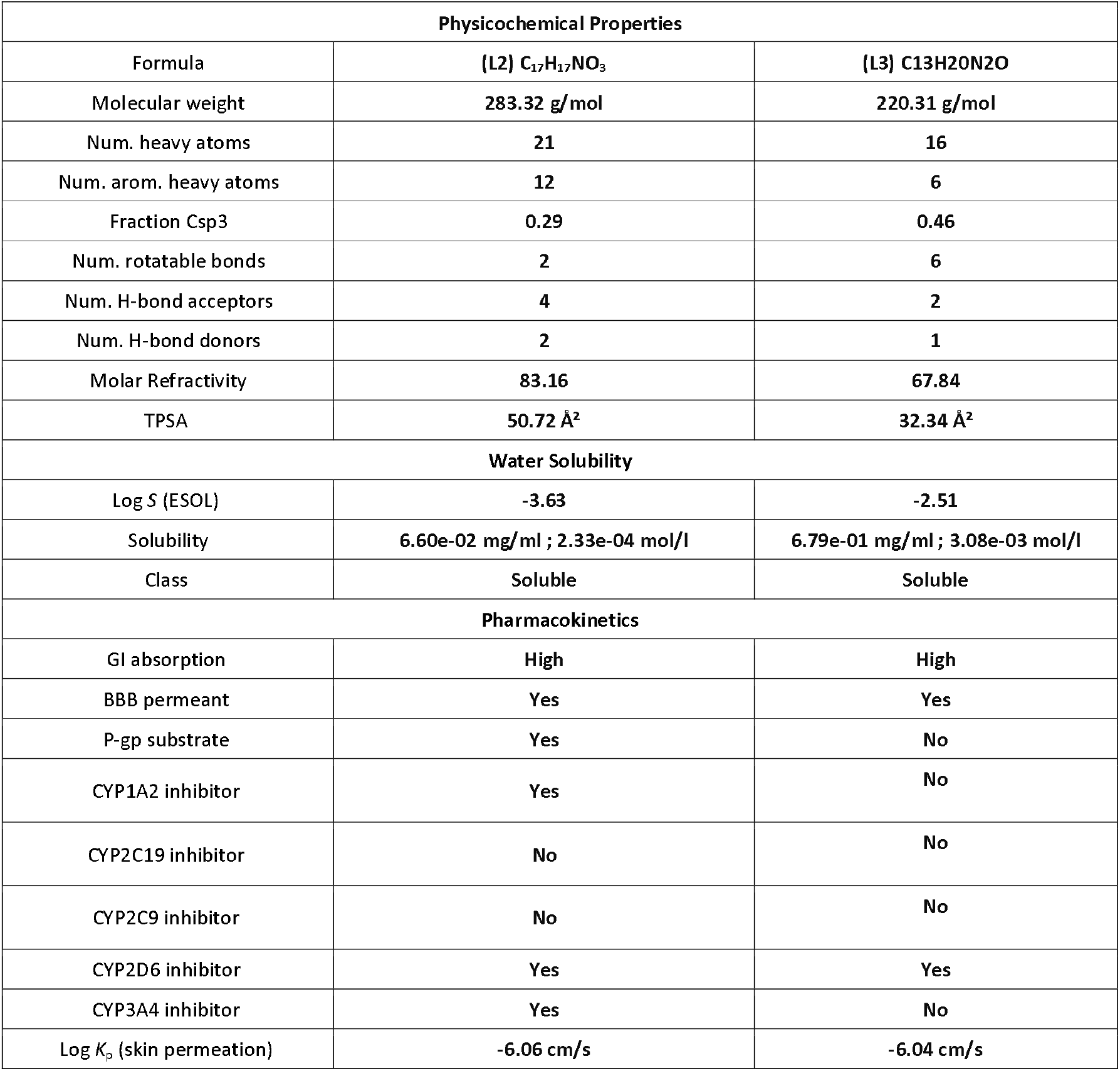
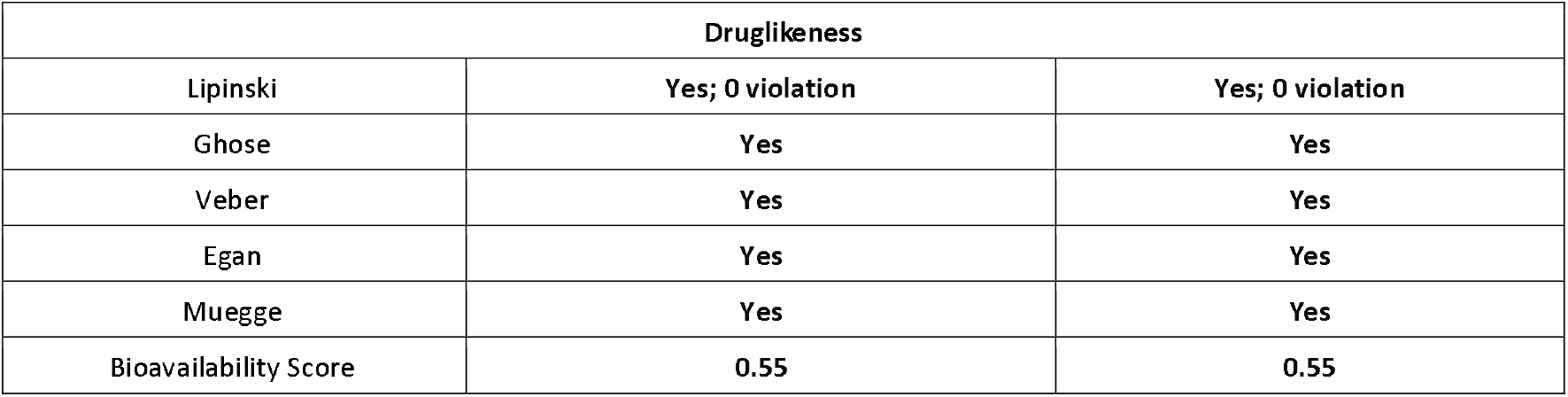
ADME analysis of 2-Diethylamino-N-P-Tolyl-Acetamide and Norcinnamolaurine.

**Table 5.**
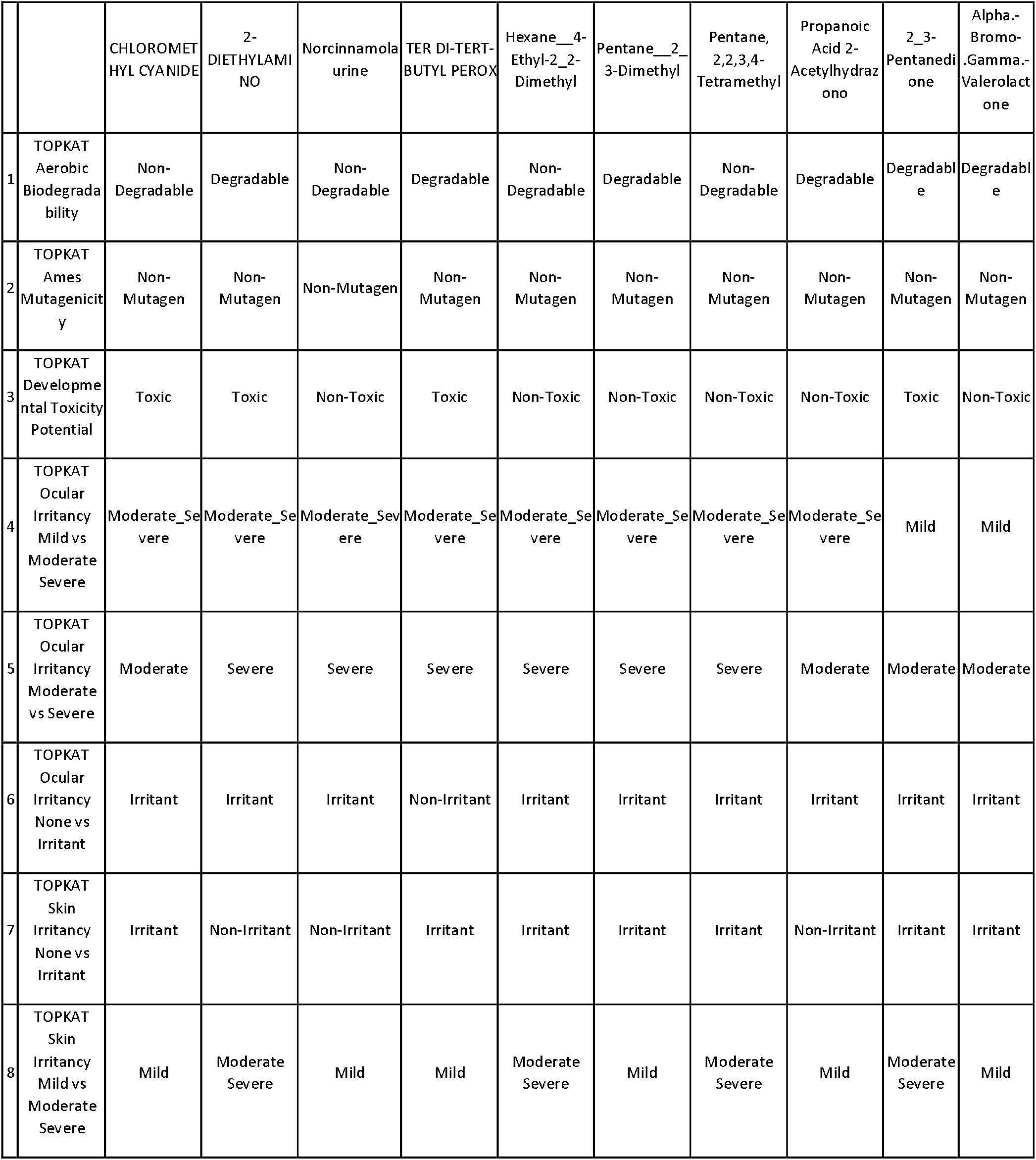
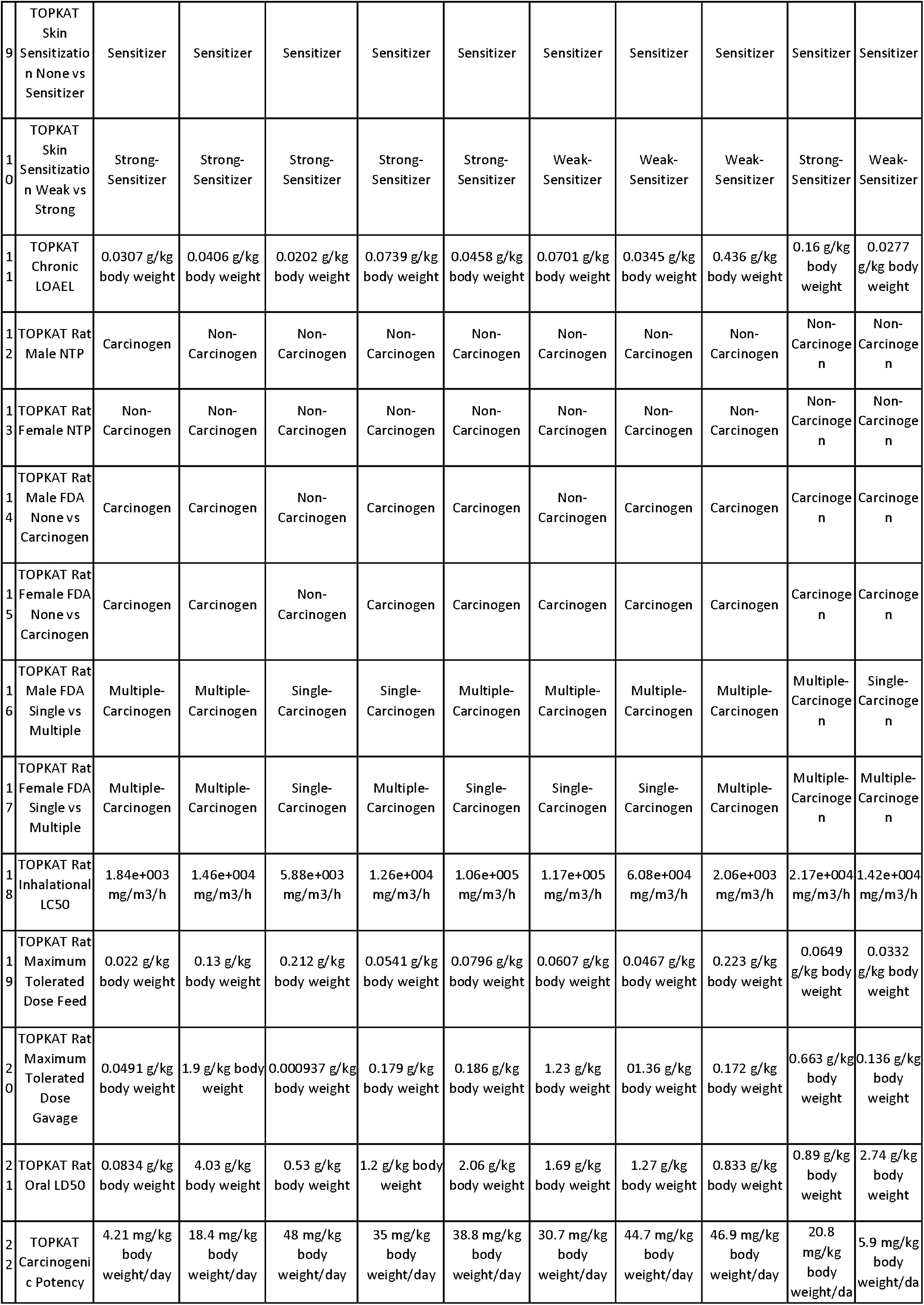
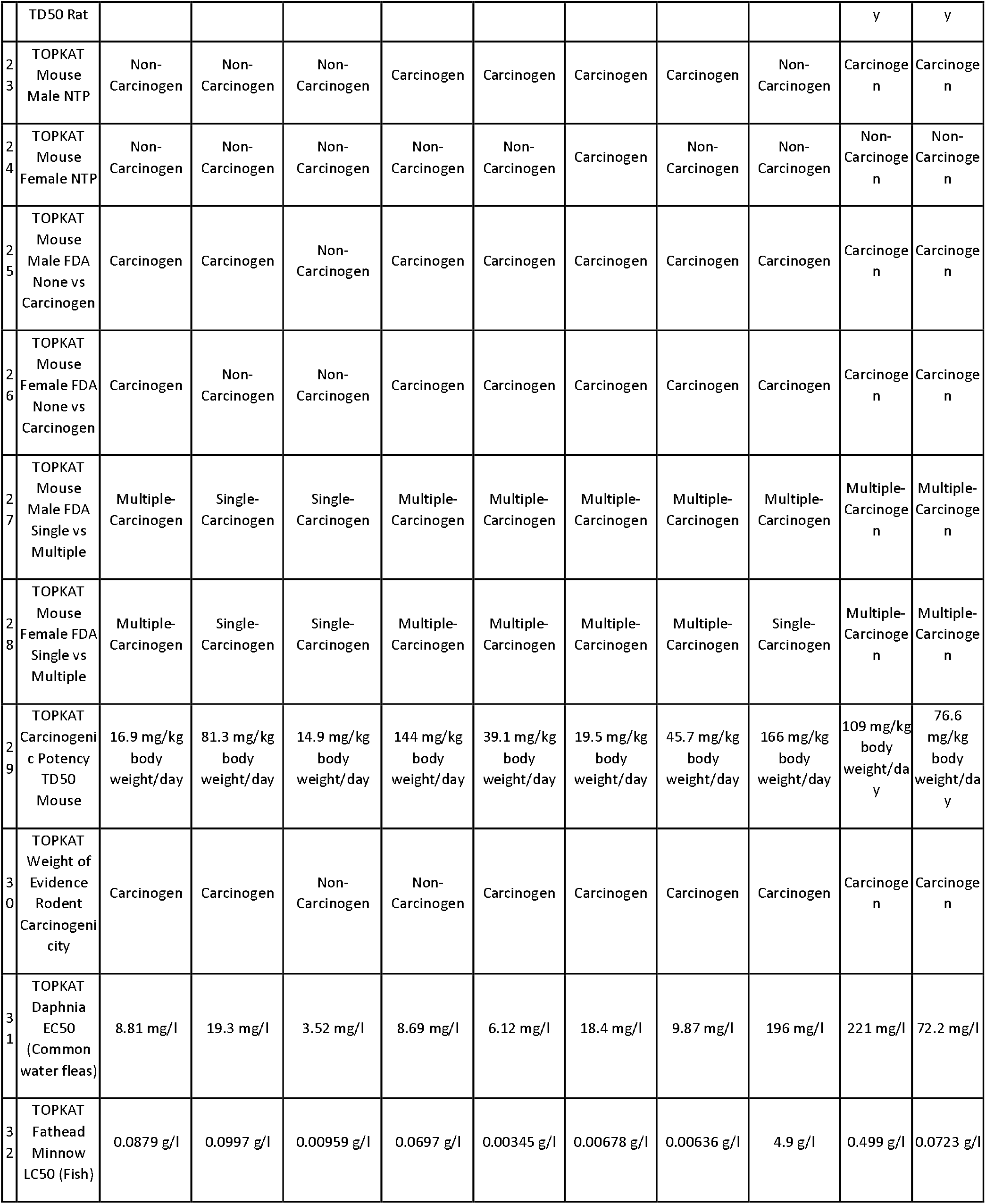
TOPKAT analysis of bioactive compounds of *E. crassipes* leaf extract.

**Table 6.**
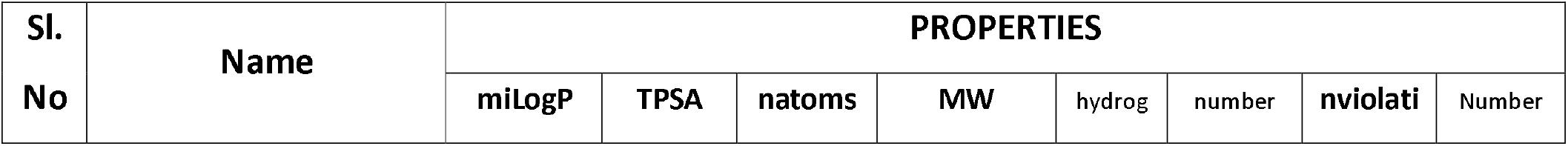
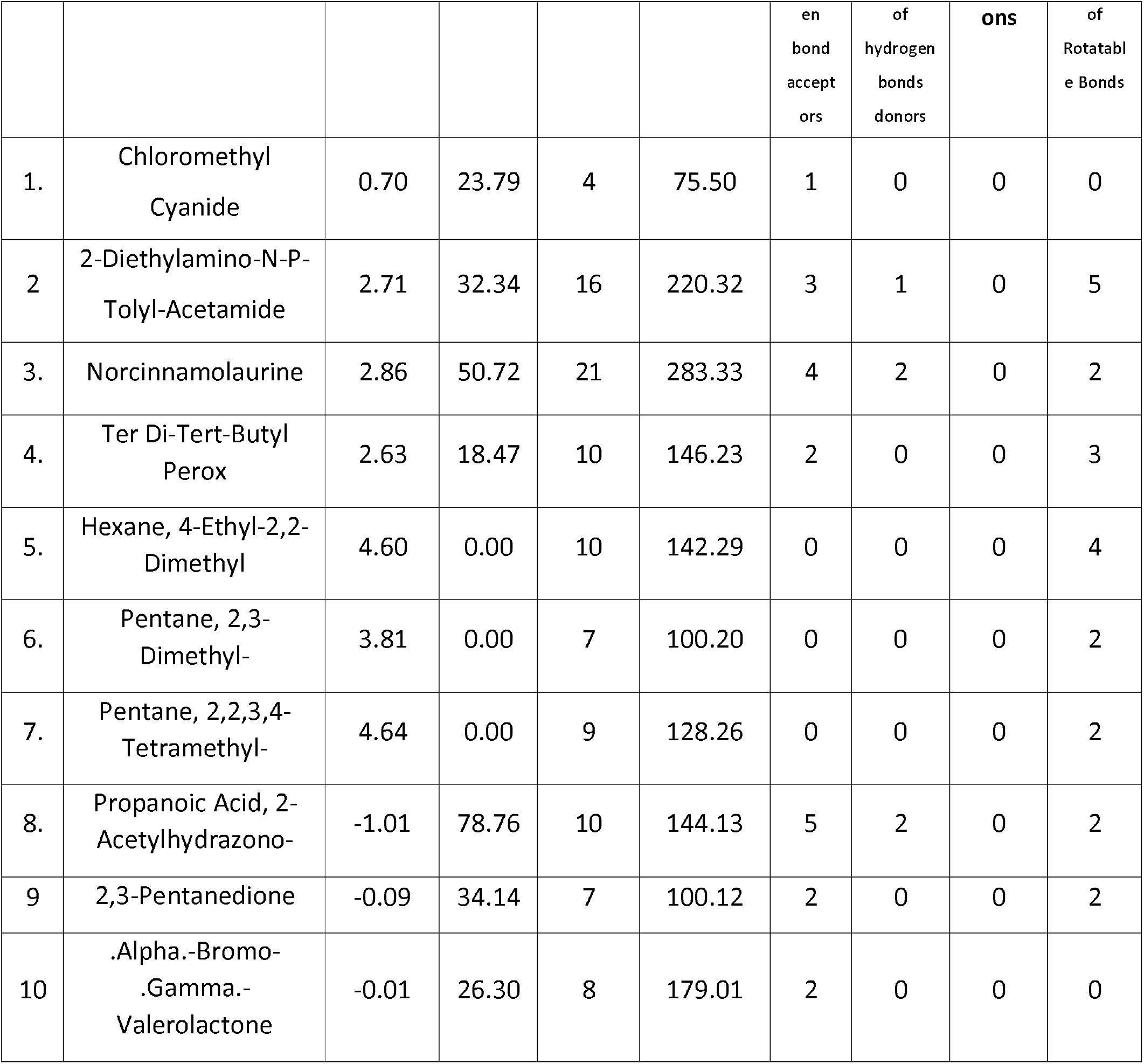
Drug-likeness scores of bioactive compounds of *E. crassipes* leaf extract.

**Table 7.**
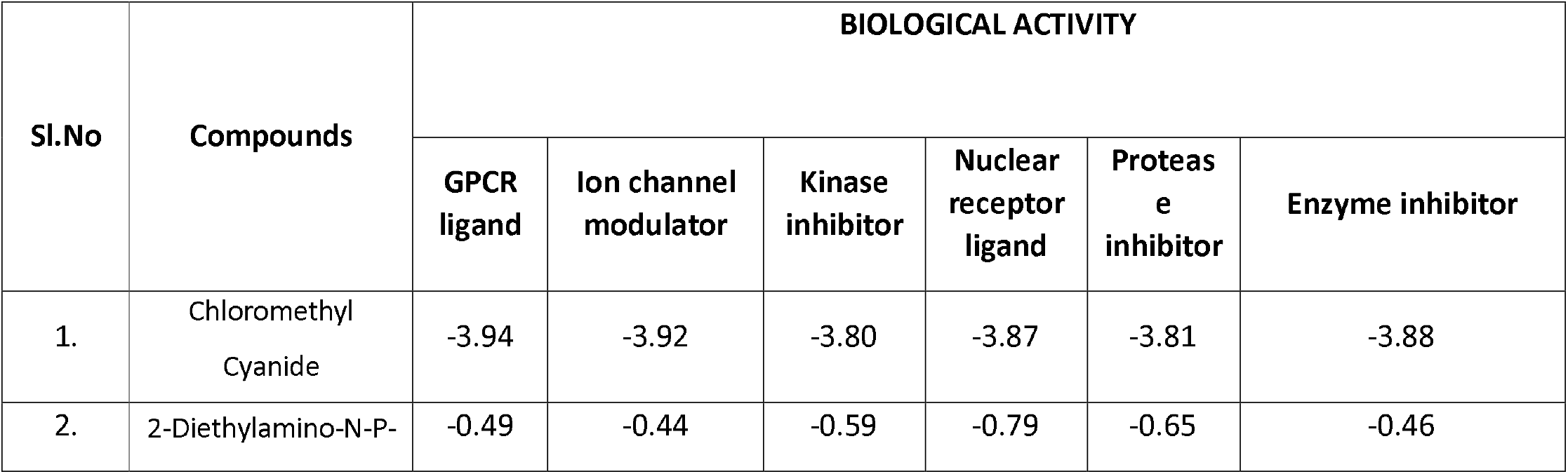
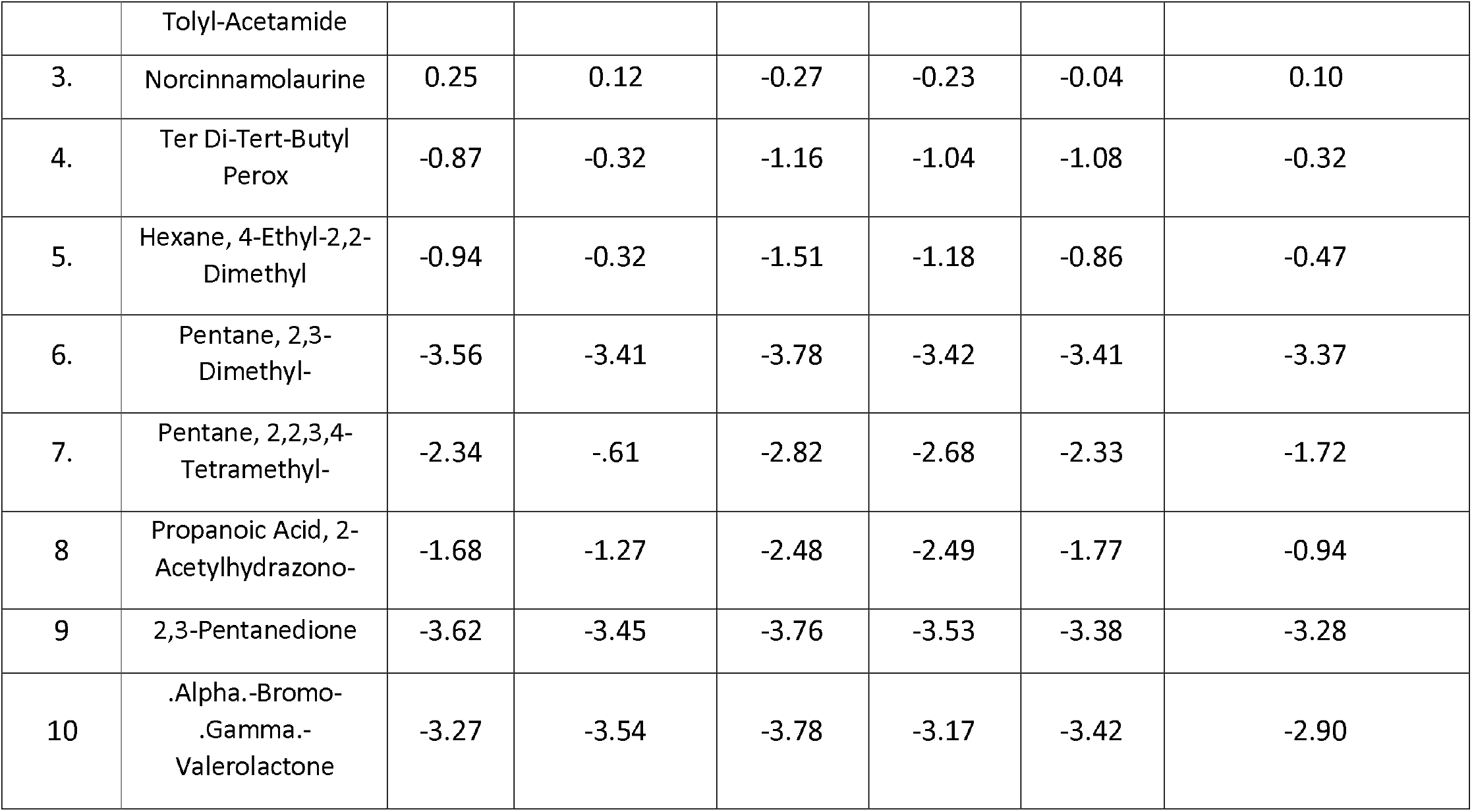
Biological activity scores of bioactive compounds of *E. crassipes* leaf extract.

